# A model of the CA1 field rhythms

**DOI:** 10.1101/2021.04.20.440634

**Authors:** Mysin I.E.

## Abstract

We propose a model of the main rhythms in the hippocampal CA1 field: theta rhythm, slow, middle, and fast gamma rhythms, and ripples oscillations. We have based this on data obtained from animals behaving freely. We have considered the modes of neuronal discharges and the occurrence of local field potential (LFP) oscillations in the theta and non-theta states at different inputs from the CA3 field, the medial entorhinal cortex, and the medial septum. In our work, we tried to reproduce the main experimental phenomena about rhythms in the CA1 field: the coupling of neurons to the phase of rhythms, cross-rhythm phase-phase and phase-amplitude coupling. Using computational experiments, we have proved the hypothesis that the descending phase of the theta rhythm in the CA1 field is formed by the input from the CA3 field via the Shaffer collaterals, and the ascending phase of the theta rhythm is formed by the inhibitory postsynaptic potentials from CCK basket cells. The slow gamma rhythm is coupled to the descending phase of the theta rhythm, since it also depends on the arrival of the signal via the Shaffer collaterals. The middle gamma rhythm is formed by the excitatory postsynaptic potentials of the principal neurons of the third layer of the entorhinal cortex, corresponds to experimental data. We were able to unite in a single mathematical model several theoretical ideas about the mechanisms of rhythmic processes in the CA1 field of the hippocampus.

## Introduction

Despite a century of historical research on the brain’s electrographic rhythms, the mechanisms of rhythm generation and their role in brain function remain one of the puzzles of modern neuroscience. In this paper, we propose a theoretical model of the mechanisms of the generation of basic rhythms in the CA1 field of the hippocampus. We chose the hippocampal CA1 field as the object for modeling for two reasons. First, the hippocampus plays a critical role in attention and memory processes (Buzsáki & Moser, 2013; Vinogradova, 2001). Second, among all hippocampal regions, the CA1 field is the most studied in the context of rhythms (Buzsáki & Wang, 2012; Buzsáki, 2002, 2015). The main goal of our work is to offer a single model of the main rhythms of the hippocampus - theta (4-12 Hz), slow (30-50 Hz), middle (50-90 Hz), and fast (90-150 Hz) gamma rhythms and ripple oscillations (110-200 Hz).

Based on the literature data, we believe that the main experimental facts that the model of the neural network of the CA1 field should reproduce and explain are as follows:

1. The phase relations of neuronal activity and rhythms. At the peak of the theta cycle, axo-axonal and neurogliaform neurons discharged. PV basket cells fired at the descending phase of the theta cyle. At the tought of the theta cycle, pyramidal, OLM, and bistratifired neurons have maximum firing rate. In the ascending phase of the theta cycle, ivy and CCK basket cells are discharged (Somogyi, Katona, Klausberger, Lasztóczi, & Viney, 2014; Somogyi & Klausberger, 2005).
2. Pyramidal neurons during the theta rhythm out of the place field fire infrequently, but the potential varies at the theta frequency (Csicsvari, Hirase, Czurko, & Buzsáki, 1998; Ylinen et al., 1995).
3. The phase precession of place cells relative to the theta rhythm (J O’Keefe & Recce, 1993; Skaggs, McNaughton, Wilson, & Barnes, 1996).
4. Phase coupling of the theta and gamma rhythms. The slow gamma rhythm is most pronounced at the descending phase of the theta rhythm, the middle at the maximum of the theta rhythm, and the fast gamma rhythm at the minimum of the theta rhythm. In addition, there is a constant phase difference between theta and gamma rhythms (Belluscio, Mizuseki, Schmidt, Kempter, & Buzsáki, 2012; Colgin & Moser, 2010; Colgin, 2015, 2016).
5. Antagonistic relations between ripple oscillations and theta rhythm, i.e. the absence of ripple oscillations with a powerful theta rhythm (Buzsáki, Leung, & Vanderwolf, 1983; Vandecasteele et al., 2014).
6. Parameters of ripple oscillations: the form of field potential fluctuations during a ripple event, the activity of neurons of different populations during ripple oscillations (Buzsáki, 2015; Somogyi et al., 2014).
7. The role of external inputs in the CA1 field. Theta rhythm generation requires input from the medial septum (MS) (Colgin, 2013; Vinogradova, 1995), slow gamma rhythm and ripple oscillations depend on input via Shaffer collaterals from the CA3 field, and the middle gamma rhythm depends on input via the perforant path from cells of the entorhinal cortex layer 3 (MEC) (Belluscio et al., 2012; Colgin et al., 2009; Colgin, 2015; Fernández-Ruiz et al., 2017).

Many mathematical models have been proposed in the literature that describes many of the presented experimental observations individually (Börgers, Talei Franzesi, Lebeau, Boyden, & Kopell, 2012; Burgess & O’Keefe, 2011; Chance, 2012; Keeley, Fenton, & Rinzel, 2017; Mysin, Kitchigina, & Kazanovich, 2019; Saraga, Ng, & Skinner, 2006; Wang & Buzsáki, 1996). However, there is no model describing all observations in a single model with a single set of parameters. In our work, we tried to combine theoretical ideas about the mechanisms of rhythms, as well as express some of our ideas about how rhythms are generated.

In experimental studies of the properties of rhythms, more attention is usually paid to the dorsal hippocampus, so our model describes the behavior of the CA1 field network of the dorsal hippocampus.

## Methods

Our model included pyramidal neurons, as well as 8 types of interneurons: CCK and PV basket cells, axo-axonal cells, bistratified cells, OLM cells, neurogliaform cells, ivy cells, and Schaffer collaterals-associated cells (Fig. 1). To describe the neuron models, we used the Hodgkin-Huxley formalism. All the neurons were multicomponent. Neuron models were taken from articles (Bezaire, Raikov, Burk, Vyas, & Soltesz, 2016; Cutsuridis & Poirazi, 2015). The connections between the neurons were taken based on the database “hippocampome” (Tecuatl, Wheeler, Sutton, & Ascoli, 2020), GABA-A receptors were modeled - for inhibitory synapses, nicotine receptors, and AMPAR for excitatory connections. Gap junctions between PV basket cells and between neurogiaform cells were included in the model in accordance with the experimental data (Fukuda & Kosaka, 2000, 2003; Price et al., 2005).

**Figure 1.**
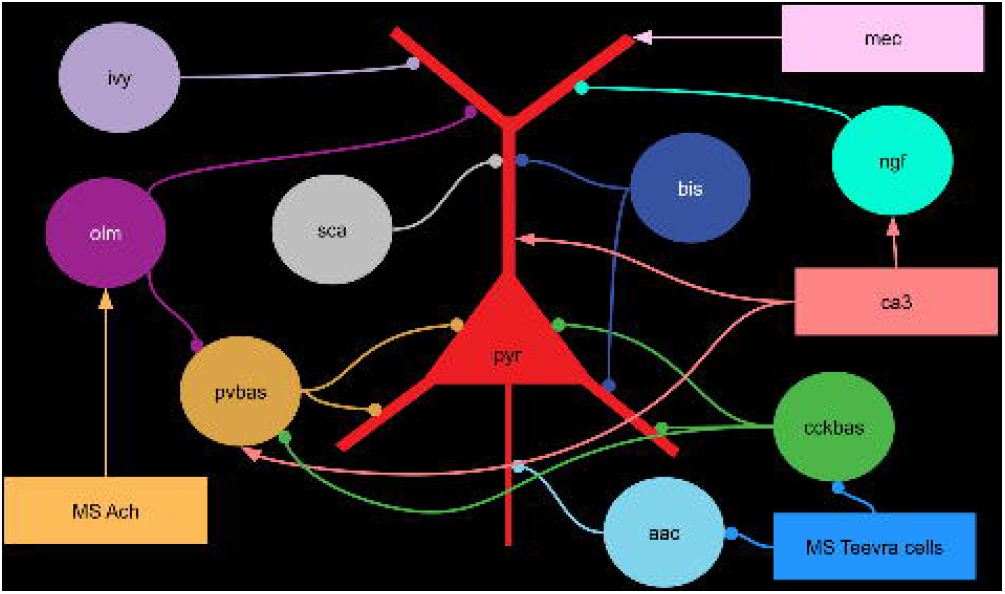
Scheme of the connections in the model. Neuron notation: pyr is pyramidal cells, pvbas is PV basket cells, olm is OLM cells, cckbas is CCK basket cells, ivy is Ivy cells, ngf is neurogliaform cells, bis is bistratified cells, aac is axo-axonic cells, sca is Schaffer collateral-associated cells. Although the interneurons are represented by circles, they were modeled by multicompartment models, see supplemental material. Not all connections are shown in the figure.

We have simulated external inputs from the CA3 field, the third layer of the entorhinal cortex, and the GABAergic and cholinergic input from the medial septum (MS). The inputs from the CA3 field were divided into spatially modulated and spatially non-modulated. The first ones imitated the activity of place cells in the CA3 field, having a peak of discharges in time, these generators showed modulation by theta rhythm (7 Hz), slow gamma rhythm (35 Hz). Spatially unmodulated generators of the CA3 field imitated the background activity of neurons of this field, showing modulation only by the theta rhythm. Generators simulating the input from the MS with a different phase shift showed similar activity. The MEC generators were modeled as grid cells, with slow periodic activity. They were also modulated by the theta rhythm and the middle gamma rhythm. The mathematical apparatus for describing spike generators and their parameters are described in the supplemental material.

To simulate the non-theta state, we removed the MEC input, spatially unmodulated CA3 generators, and the cholinergic input from the MS from the model. The Teevra cells went into a constant discharge mode (Viney et al., 2013). Spatially modulated generators were modulated by a slow gamma rhythm (35 Hz), ripple oscillations (170 Hz), and the centers of their discharges converged.

The weights of connections from spatially modulated CA3 generators to pyramidal and PV basket neurons were set using the Gaussian function. The weights of the connections from the MEC generators to the pyramidal neurons were also distributed using the Gaussian function, but the center for the pyramidal neurons was shifted by 500 ms to the beginning of the simulation. PV basket neurons, in turn, inhibited mainly distant pyramidal neurons, the experimental literature provides indirect evidence of such a distribution of connections (Hangya, Borhegyi, Szilágyi, Freund, & Varga, 2009; Udakis, Pedrosa, Chamberlain, Clopath, & Mellor, 2020). All other connections between neurons in our model were established randomly, the density and other parameters are given in the supplement.

The LFP has been simulated using linear approximation (Parasuram et al., 2016). The simulations were performed in the Neuron simulator. For parameters, see the supplemental materials. The data analysis was carried out in a similar way to the experimental work. The methods of analysis are also given in the supplement.

## Results

### General characteristics of the model

In our work, we tried to combine many theoretical ideas about how the main rhythms are formed in the CA1 field of the hippocampus. From a practical point of view, the purpose was to optimize the parameters of the model, primarily the weights of connections between populations of neurons, in such a way as to harmonize theoretical ideas about the formation of different rhythms. In the process of optimizing the parameters of the model, two tasks can be distinguished: the explanation of the shape of the field potential and the dynamics of the discharges of neurons during rhythmic processes.

According to modern concepts, the field potential is formed by currents in the extracellular environment, which are produced by a currents through the membranes of pyramidal neurons. The currents through the membranes of pyramidal neurons are summed up due to the ordered structure of the processes of pyramidal neurons (Buzsáki, Anastassiou, & Koch, 2012; Parasuram et al., 2016). Based on the analysis of LFP signals by the current source density method, it is possible to identify which layers are involved in the generation of each rhythm. When constructing our model, we optimized the parameters so that the inputs from the CA1 interneurons and external inputs create the corresponding currents and provide depolarization or hyperpolarization of the corresponding compartments of the pyramidal neurons. For example, with the slow gamma rhythm, the maximum currents occur between the stratum radiatum and the stratum oriens, which corresponds to the excitation of pyramidal neurons by Shaffer collaterals and the inhibition by PV basket cells, so the connections from these neurons to the pyramidal neurons in our model are large.

In the CA1 field, almost all neural populations are modulated by rhythms. The problem of forming the dynamics of neuronal activity observed in the experiment was solved by optimizing the parameters of connections from external inputs and between neurons in such a way that they formed the experimentally observed dynamics relative to each other. For example, during the theta rhythm of PV and CCK basket neurons are discharged in opposite phases, descending and ascending, respectively. In our model, we established large weights of connections between them. Thus, antiphase relationships are formed between these neural populations.

Fig. 2, 3, 4 show the results for the theta state, and Fig. 8 shows the results for the non-theta state. These results correspond to the experimental data listed in the introduction. In the following sections, we describe in detail how we obtained these results.

**Figure 2.**
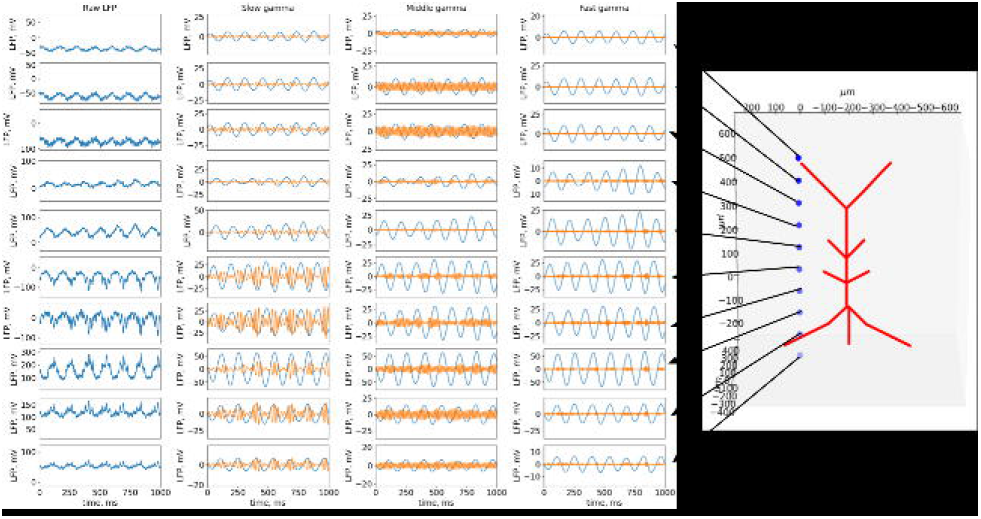
Raw LFP and oscilation bands dynamics across layers in the theta state. The morphology of the pyramidal neuron is shown on the right. The blue dots indicate the sites where the LFP was calculated.

**Figure 3.**
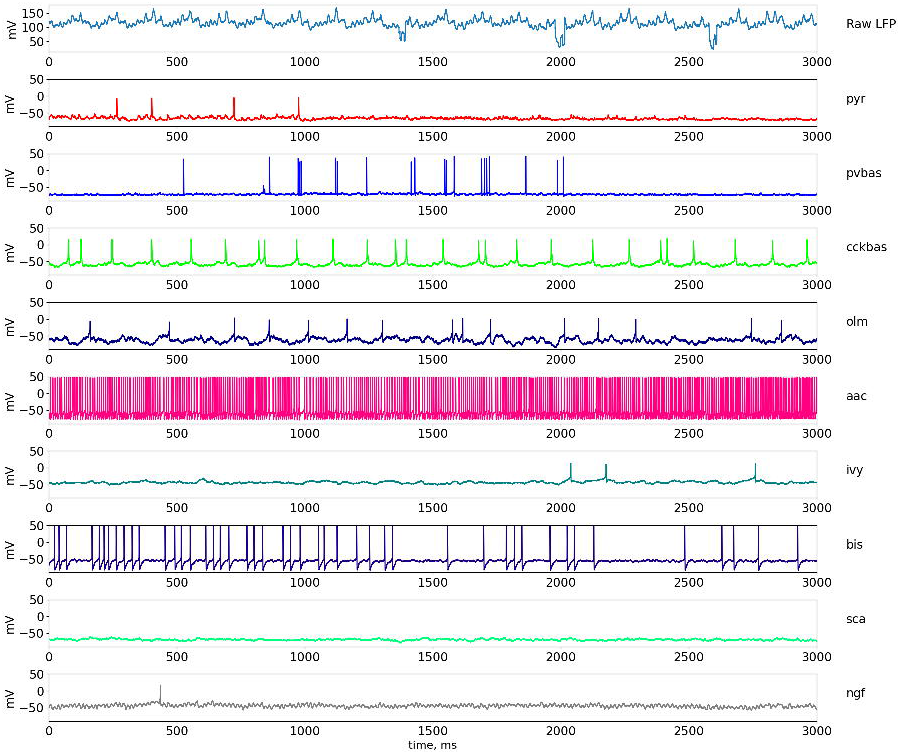
Dynamics of LFP and neuronal activity in the theta state. The upper part of the panel shows the simulated LFP in the pyramid layer. Examples of the dynamics of intracellular potentials on the soma of neurons are also presented below. The notation of neurons are similar to Fig. 1.

**Figure 4.**
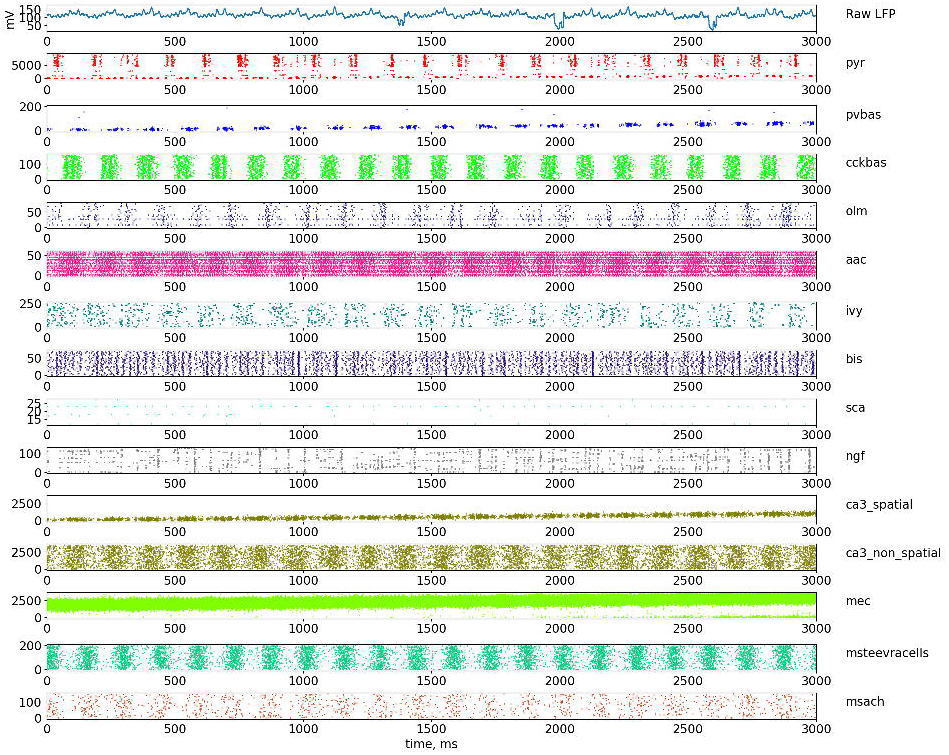
Dynamics of LFP and neuronal firing in the theta state. The upper part of the panel shows the simulated LFP in the pyramid layer. The lowest series of plots show raster graphs of the discharges of neurons. The notation of neurons are similar to Fig. 1.

### Model of theta rhythm

The theoretical ideas about the mechanisms of theta rhythm suggested in this paper are based on our previous work (Mysin et al., 2019). In our previous study, we showed that input from the CA3 field via Shaffer collaterals plays a key role in theta rhythm generation in the CA1 field. This hypothesis explains the biological plausibility of the activity of pyramidal and PV basket neurons during the theta cycle. In this work, other types of interneurons were have added to our model, in particular CCK basket cells. We believe that CCK basket neurons are also important for theta rhythm generation. The discharges of these cells occur in the ascending phase of the theta rhythm in the pyramidal layer of the CA1 field (Fig. 5)(Somogyi & Klausberger, 2005), which coincides with the hyperpolarization of the soma of pyramidal neurons. In addition, CCK basket neurons receive very powerful input from the medial septum (Joshi, Salib, Viney, Dupret, & Somogyi, 2017). Thus, in this model, the theta cycle is formed by two mechanisms: depolarization of the perisomatic zone of pyramidal neurons by the input from Shaffer collaterals in the descending phase of the theta rhythm and hyperpolarization of the soma of pyramidal neurons by the inputs from CCK basket neurons. The rhythmic activity of Shaffer collaterals is embedded in the model. The rhythmicity of the CCK basket neurons occurs due to the presence of two inputs from the medial septum - cholinergic and GABAergic (Joshi et al., 2017). The cholinergic input has a muscarinic nature and is modeled by the presence of a tonic excitatory current in the CCK basket neurons. GABAergic input is provided by rhythmic spike generators that mimic the input from the Teevra cells of the medial septum (Joshi et al., 2017). Due to these two mechanisms, CCK basket cells are introduced into a stable rhythmic mode.

**Figure 5.**
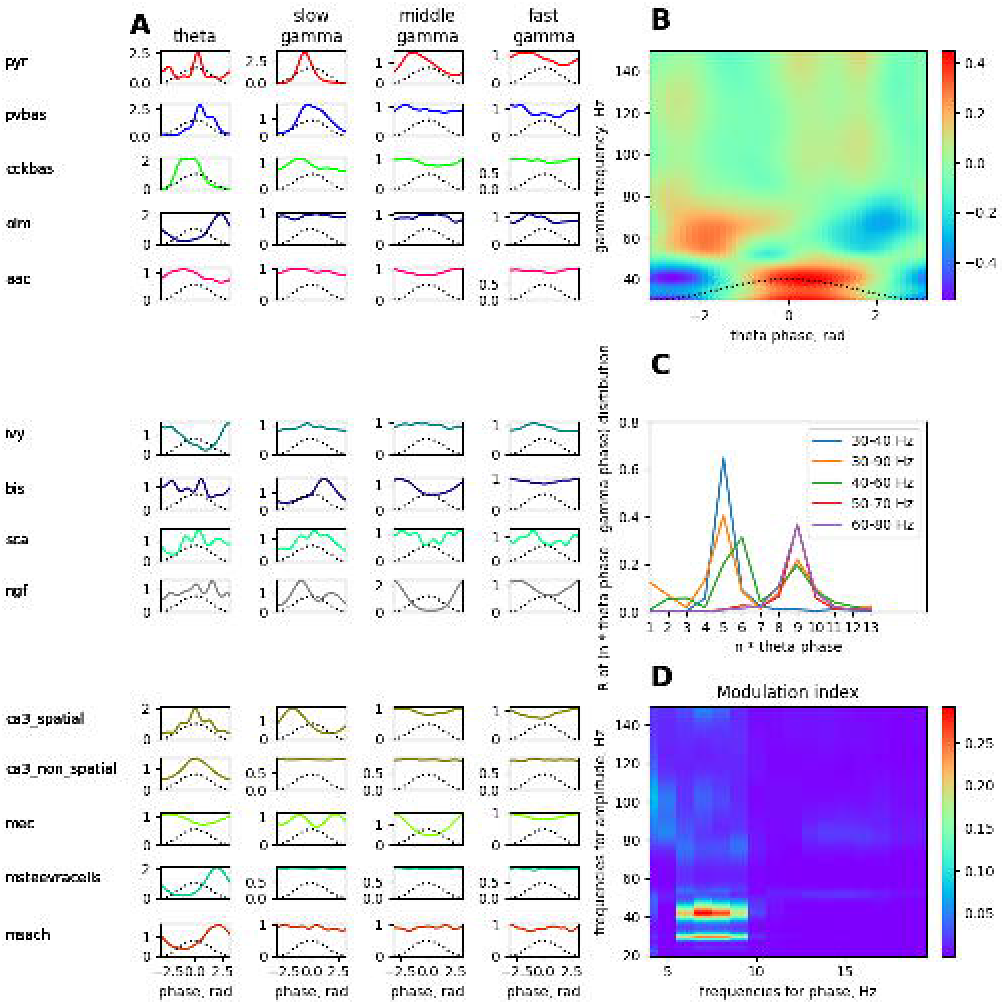
Phase analysis of the dynamics of neurons relative to LFP rhythms and cross-rhythm relations in the theta state. A simulated LFP signal from the pyramid layer was used to construct all the plots. **A**. Distribution of spikes by rhythm phases. **B**. Amplitude-phase coupling of gamma rhythms and theta rhythms. **C**. Phase-phase coupling of theta and gamma rhythms n:m-test. **D**. The modulation index of the amplitude from the range of the gamma rhythm by the phase of the theta rhythm. The notation of neurons are similar to Fig. 1.

**Figure 6.**
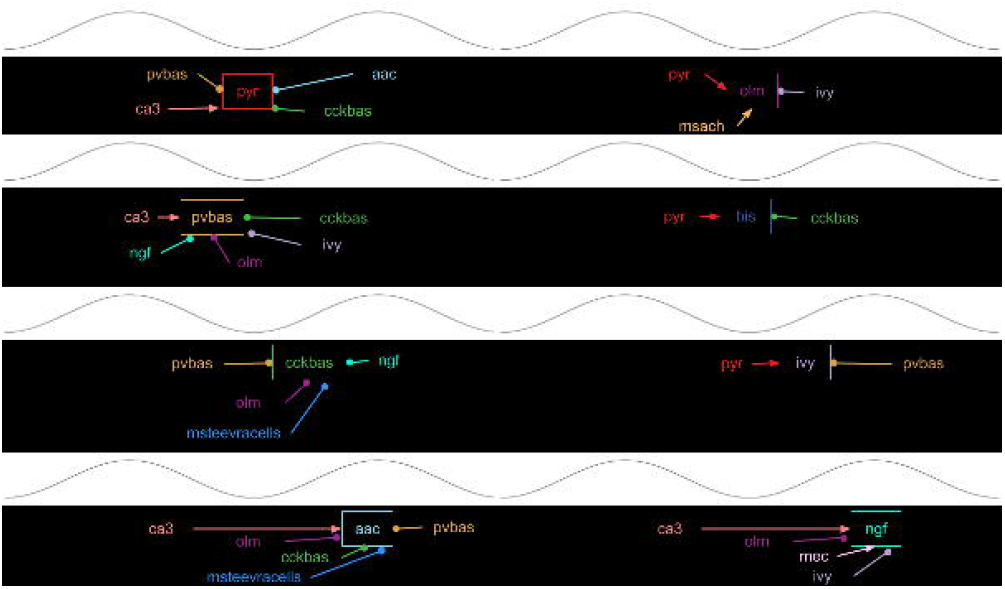
The scheme of coupling between neuronal population firing and the theta rhythm phase. The notation of neurons are similar to Fig. 1.

The role of interneurons, except for the CCK basket neurons, in the generation of theta rhythm, is currently unclear. Available experimental data on PV basket and OLM neurons indicate that these cells do not play a critical role in the generation of theta rhythm in the CA1 field (Royer et al., 2012). We found no experimental data on the functions of the other neurons. In our model, we optimized the parameters of connections from other interneurons, except for the CCK basket neurons, so that their inputs to the pyramidal neurons do not destroy the theta rhythm mechanism described above. The connections between the populations of interneurons were optimized so that they formed the correct coupling of the discharges to the theta rhythm phase (Fig. 5).

It should be noted that the distribution of firing of pyramidal neurons in the theta rhythm phase is bimodal in our model (Fig. 5A). Pyramidal neurons outside the place field are essentially discharged at the minimum of the theta rhythm. However, inside the place field, the inhibition of PV basket cells is less (Fig. 7C), so the pyramidal neurons begin to discharge at the descending phase of the theta rhythm. In our model, the ratio of spatially modulated and unmodulated pyramidal neurons is 1:1, which creates two pronounced peaks on the theta rhythm phase distribution plot. The discharges of the place cells in the descending phase of the theta rhythm in our model play a phase role in the formation of a slow gamma rhythm, which we will discuss below. We could achieve a single dominant peak at the theta rhythm minimum by increasing the number of spatially unmodulated pyramidal neurons, but this would increase the computational complexity of the model.

**Figure 7.**
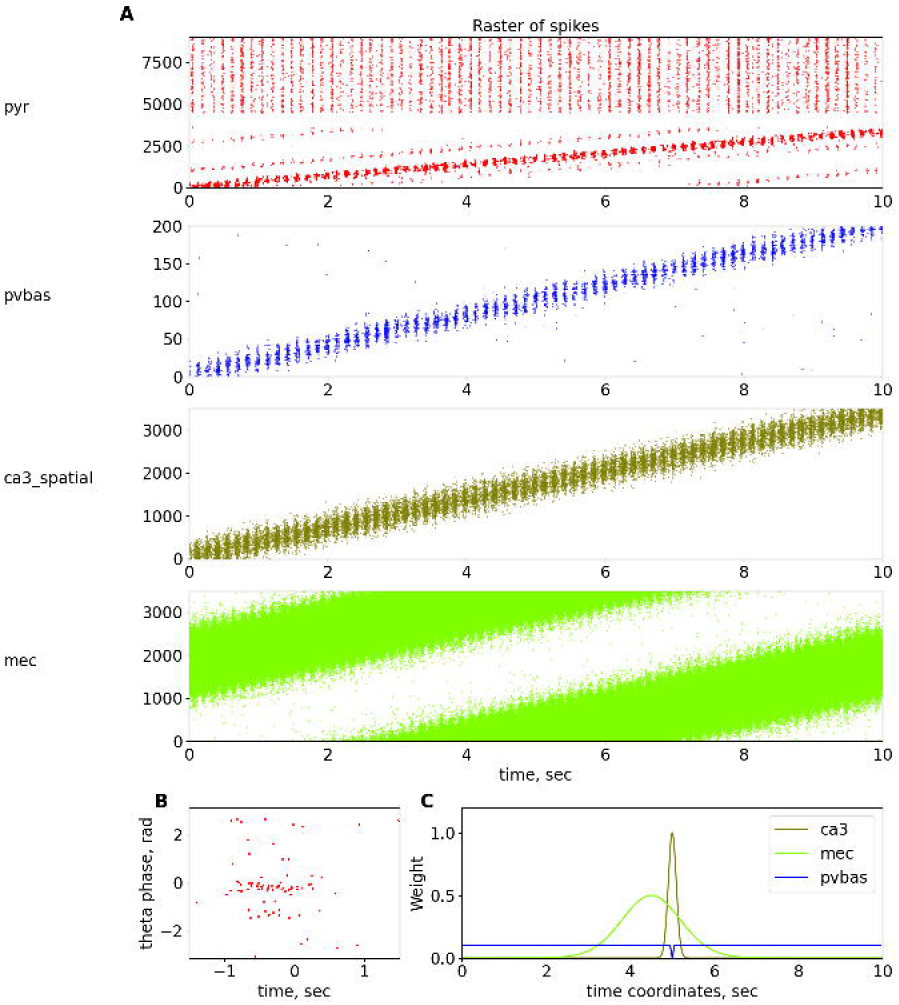
Place cells. **A**. Raster plots of spatially modulated inputs and model neurons. **B**. Phase precession of pyramidal neurons. **C**. Dependence of weights of connetion from ca3_spatial, mec and pvbas to the pyramydal cells from their coordinates, for detail see supplement material (Table 22). The notation of neurons are similar to Fig. 1.

### Place cells and phase precession

In our work, we focused on the mechanisms of rhythms, but we could not avoid the topic of place cells, because the phenomenon of place cells is closely related to the mechanisms of rhythm generation. In our model, we will not consider the processes of learning, place cell formation, and remapping. The place cells in our model are present only to the extent that we believe is necessary to investigate the mechanisms of the rhythms. The activity of pyramidal neurons as place cells is inherited from inputs from the CA3 field and the entorhinal cortex (Fig. 7). It is shown that pyramidal neurons are modulated by slow gamma rhythm, when they are discharged the animal is inside the place field of these neurons (Senior, Huxter, Allen, O’Neill, & Csicsvari, 2008), this will be discussed in more detail later in the section on gamma rhythm.

Another important physiological phenomenon that links rhythmic field activity and cell discharges is phase precession (Burgess & O’Keefe, 2011; J O’Keefe & Recce, 1993). A large number of theoretical explanations of the phase precession phenomenon have been proposed, although most hypotheses are reduced to two theoretical ideas: the interference of two-oscillator inputs and the asymmetry of one exciting input (Burgess & O’Keefe, 2011). In our work, we will adhere to the hypothesis based on the interference of two inputs: from the CA3 field and the entorhinal cortex (Chance, 2012; Fernández-Ruiz et al., 2017). In our model, we optimized the parameters, focusing primarily on the data of the article (Fernández-Ruiz et al., 2017), because we believe this hypothesis is the most reasonable. In our model, the input from the entorhinal cortex comes earlier than the input from the Shaffer collaterals, which ensures the beginning of the discharges of the place cells first from one input and then from another (Fig. 7), which ultimately allows us to reproduce the dependence of the theta rhythm phase on the position of the animal for the discharges of the place cells. It should be noted that in our model, the phase precession is not as pronounced as in the special models devoted to this phenomenon. This is due to the fact that our model contains additional elements, in particular CCK interneurons. The presence of many interneurons leads to the fact that pyramidal neurons receive more rhythmic inputs, in contrast to many other models. We could not optimize the parameters of our model for a better description of the phase precession without reducing the quality of the description of the properties of rhythms. We believe that an experimental study of the role of different groups of interneurons in the formation of place cells should provide the necessary data to improve a model of phase precession. We did not introduce additional hypotheses about the spatial modulation of different populations of interneurons, so as not to complicate the model.

### Model of slow gamma rhythm

Currently, it has been proved that PV basket neurons play a critical role in the generation of the slow gamma rhythm. There are two theoretical schemes to explain the mechanisms of their synchronization (Buzsáki & Wang, 2012; Colgin & Moser, 2010). The first mechanism involves the synchronization of the PV basket neurons with each other due to inhibitory synapses and gap junctions (Saraga et al., 2006; Wang & Buzsáki, 1996). The second mechanism involves the synchronization of pyramidal neurons and the excitation from them of the PV basket neurons, which in turn give inhibitory feedback to the pyramidal neurons (Börgers et al., 2012; Keeley et al., 2017). The second mechanism has the most experimental confirmation, but it does not contradict the first one. In our model, we included both mechanisms. We included gap junctions between PV basket neurons. Also, the synapses between pyramidal and PV basket neurons are optimized in such a way that inhibitory feedback between them was active.

Many studies have shown that slow gamma occurs in the descending phase of the theta rhythm in the pyramidal layer, and it is a consequence of the arrival of the signal from the Shaffer collaterals (Belluscio et al., 2012; Colgin et al., 2009; Colgin, 2015; Fernández-Ruiz et al., 2017). As we discussed in the section on the mechanisms of the theta rhythm, we believe that the descending phase of the theta rhythm is provided by the Shaffer collaterals input. The excitation from the Shaffer collaterals leads to the excitation of a part of the pyramidal neurons, from which the PV basket neurons are excited. Further, the PV neurons inhibit the pyramidal neurons, as a result, negative feedback is formed, which in the field potential manifests itself as a slow gamma rhythm. The mechanism of coupling of theta and gamma rhythms consists of a single reason for the descending phase of the theta rhythm is the arrival of a signal via the Shaffer collaterals (Fig. 4).

The presence of gap junctions between the PV basket neurons and strong connections between the pyramidal and PV basket neurons creates the possibility of forming gamma oscillations in the CA1 field. The pyramidal neurons of the CA3 field also have modulation at the gamma frequency, the rhythmic input by the Shaffer collaterals resonates with the endogenous mechanisms of gamma rhythm generation in the CA1 field, as a result, we observe high coherence at the gamma frequency between the signals registered in the CA3 and CA1 fields (Colgin, 2015; Fernández-Ruiz et al., 2017). In our model, only the spatially modulated generators that mimic the place cells in the CA3 field showed modulation by the slow gamma rhythm. This allows us to explain the experimental fact that the activity of pyramidal neurons inside their place field is more strongly modulated by a slow gamma rhythm than outside the field of place (Senior et al., 2008).

The coupling parameters of the other interneurons (except for the PV basket interneurons) were not optimized for simulating the slow gamma rhythm (Fig. 2). We did this for two reasons. On the one hand, the literature does not show the participation of other groups of interneurons in the generation of a slow gamma rhythm. On the other hand, the slow gamma rhythm is most powerful in the descending phase of the theta rhythm, at this phase of the theta rhythm, only the PV basket neurons have a peak of activity, the other interneurons are discharged little and as a result, their action potentials play a small role in the dynamics of the network.

### Model of middle gamma rhythm

Currently, theoretical models of the middle gamma rhythm are not proposed. However, the authors of the experimental articles agree that the middle gamma rhythm in the CA1 field is a consequence of the arrival of a signal from the entorhinal cortex via the perforant path, and it is also well established that the middle gamma rhythm is associated with the theta rhythm (Belluscio et al., 2012; Colgin et al., 2009; Colgin, 2015; Fernández-Ruiz et al., 2017). In our model, the input from the entorhinal cortex was emitted by artificial spike generators that were modulated at the frequency of the middle gamma rhythm (63 Hz). Thus, in our model, the mechanism of the middle gamma rhythm consists of induction by the entorhinal cortex, which corresponds to the experimental data (Fig. 2, 3, 5).

### Model of fast gamma rhythm

No theoretical models of the fast gamma rhythm have been proposed in the literature. Experimental data on this rhythmic process are scant. The maximum power of the fast gamma rhythm is observed in the pyramidal layer of the CA1 field near the minimum of the theta rhythm (Fernández-Ruiz et al., 2017), during the maximum firing of pyramidal neurons (Somogyi et al., 2014) (Fig. 2, 3, 5). These two facts allow us to hypothesize that the fast gamma rhythm is formed due to the generation of action potentials by pyramidal neurons. In our model, the mechanism of the fast gamma rhythm is based on this hypothesis.

### Model of ripples oscillations

Several models of ripple oscillations have been proposed in the literature (Jahnke, Timme, & Memmesheimer, 2015; Malerba, Krishnan, Fellous, & Bazhenov, 2016; Omura, Carvalho, Inokuchi, & Fukai, 2015; Stacey, Lazarewicz, & Litt, 2009; Taxidis, Coombes, Mason, & Owen, 2012; Taxidis, Mizuseki, Mason, & Owen, 2013), describing the signal form of the field potential during the generation of a ripple event, the dynamics of neuron activity during a ripple event, as well as the phase relationship between neuron activity and field potential (Buzsáki, 2015; Klausberger et al., 2005; Varga, Golshani, & Soltesz, 2012). Experimental data indicate that the mechanism for generating ripple oscillations is the rapid synchronization of pyramidal neurons due to excitatory connections between themselves. Such activity is more likely to occur in the dentate gyrus or the CA3 field, because there are very strong collateral connections between the principal neurons, and this activity is then projected into the CA1 field (Nakashiba, Buhl, McHugh, & Tonegawa, 2009). In our model, to simulate the transition of the hippocampal network to the non-theta state, we changed the parameters of external inputs. The input from the Shaffer collaterals became fast, and the input from the Teevra cells became constant (Joshi et al., 2017). Constant input from the Teevra cells led to constant inhibition of the CCK basket and axo-axonal neurons, which led to a decrease in the perisomatic inhibition of pyramidal neurons. Reduced perisomatic inhibition makes it possible for pyramidal neurons to discharge at a higher frequency.

In our ripple model, the event is completely formed by the input from the CA3 field. To simulate a ripple event, we reduced the discharge time of the CA3 generators by 40 times faster, while the discharge sequence time was about 100 ms (Fig. 8), which corresponds to the time parameters of ripple events (Buzsáki, 2015). The generators simulating the input from the CA3 field were modulated by the frequency of a slow gamma rhythm (35 Hz) and ripple oscillations (170 Hz), which corresponds to experimental data (Carr, Karlsson, & Frank, 2012).

**Figure 8.**
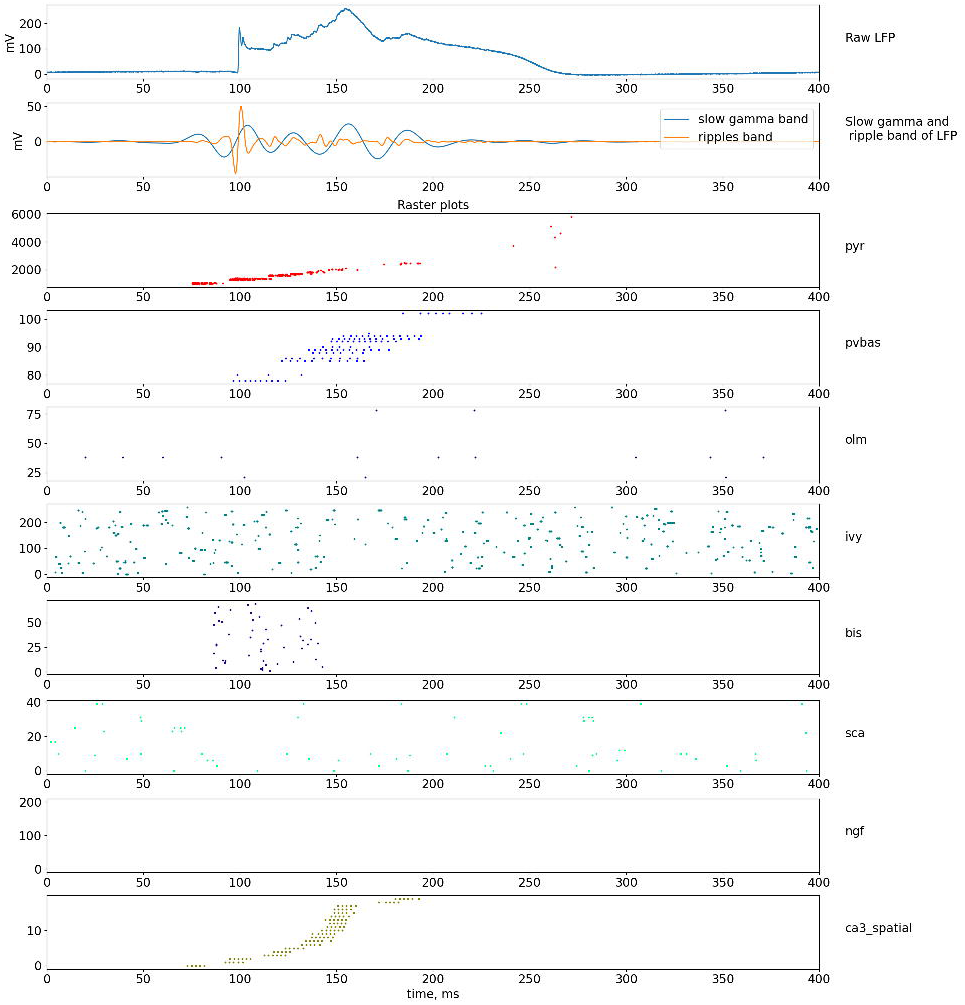
Dynamics of LFP and neuronal activity in the non-theta state. The upper part of the plots shows the simulated LFP in the pyramid layer, and the signals in the frequency bands of the slow gamma rhythm and ripple oscillations are shown below. The lowest series of plots shows raster graphs of the spikes. The notation of neurons are similar to Fig. 1.

## Discussion

### General notes

In our work, we first proposed a model of the neural network of the hippocampal CA1 field, which reproduces two main rhythmic states: theta and non-theta state, depending on the parameters of external inputs. Our model describes the role of pyramidal and eight types of interneurons in the generation of rhythmic processes of the CA1 field. One of our main goals was to reproduce and explain the dynamics of field potential fluctuations recorded *in vivo*. Therefore, in our work, we tried to take into account in detail the external inputs to the CA1 field from the smedial septum, the CA3 field, and the entorhinal cortex. Unlike many theoretical works, we did not try to get a rhythmic mode inside our model, almost all rhythms in our model are induced from external inputs. On the one hand, this corresponds to the experimental data obtained *in vivo*. On the other hand, the question of the emergence of synchronous modes in neural networks is studied in detail in theoretical works. The occurrence of theta synchronous activity in the MS is considered in several papers (Mysin, Kitchigina, & Kazanovich, 2015; Ujfalussy & Kiss, 2006). The occurrence of slow gamma rhythm and ripple oscillations have also been considered in numerous studies (Börgers et al., 2012; Jahnke et al., 2015; Keeley et al., 2017; Malerba et al., 2016; Omura et al., 2015; Saraga et al., 2006; Taxidis et al., 2012, 2013; Wang & Buzsáki, 1996). We tried to combine already known theoretical concepts in a single model and obtain a model that describes the largest possible set of experimental data on rhythms in the CA1 field. As a result, we were able to make a single model that describes all the experimental data listed in the introduction.

### Theta rhythm model

Our contribution to the proposed general model was to develop a theta rhythm model (Mysin et al., 2019). In this section, we want to discuss in detail the proposed mechanism of theta rhythm. Unlike other rhythms, we completely ignored the theta rhythm models proposed by other authors (Bezaire et al., 2016; Ferguson et al., 2015; Neymotin et al., 2013; Orbán, Kiss, & Erdi, 2006). This is due to the fact that the proposed theta rhythm models contradict a lot of important experimental data. In particular, many of the models cited to explain the occurrence of theta rhythm as a synchronization process that occurs locally in the hippocampus. However, the overwhelming amount of experimental data suggests that synchronous theta activity of neurons occurs in the medial septum, then due to GABAergic projections, it is transferred to the hippocampus (Buzsáki, 2002; Colgin, 2013; Vertes & Kocsis, 1997; Vinogradova, 1995). Theta frequency oscillations can occur in the hippocampus *in vitro* experiments under the influence of strong excitatory agents, such as carbacholine (Kazmierska & Konopacki, 2013; Traub, Bibbig, LeBeau, Buhl, & Whittington, 2004; Williams & Kauer, 1997). However, this activity is very different from the theta rhythm, which is observed in animals in free behavior. The main difference is the number of simultaneously active pyramidal neurons. When generating a theta rhythm in animals in free behavior, only about 1% of CA1 pyramidal neurons are discharged in a single theta cycle (Ylinen et al., 1995), similar experimental estimates exist for the CA3 field and the dentate gyrus (Soltesz, Bourassa, & Deschênes, 1993; Soltesz & Deschênes, 1993). In models based on the idea of synchronization within the hippocampus, the number of simultaneously active pyramidal neurons is very large and close to 100%, since synchronization occurs due to excitatory connections between the principal neurons.

The main ideas about the mechanism of theta rhythm generation are based on the results of our previous article (Mysin et al., 2019), but we have introduced additions to our theoretical scheme, which should be discussed in detail. One of the main results of our previous article was the idea of Shaffer collaterals as the main input forming the descending phase of the theta rhythm in the CA1 field. Subsequent experimental research using optogenetics methods showed our correctness. When the PV basket neurons were stimulated in the CA3 field, the theta rhythm power in the CA1 field decreased (López-Madrona et al., 2020). Our prediction about the role of the perforant path in theta rhythm generation, made in our previous work (Mysin et al., 2019), was also confirmed. The perforant path input does not play a role in theta rhythm generation, as confirmed by experiments with signal suppression from the entorhinal cortex (López-Madrona et al., 2020). Although the results (López-Madrona et al., 2020) directly contradict the results of previous studies (Middleton & McHugh, 2016). We believe that the difference lies in the methods of research. The article used a genetic modification of CA3 pyramidal neurons (Middleton & McHugh, 2016), while the article (López-Madrona et al., 2020) used optogenetic methods, the effect was acute, without adaptive changes. We believe that the data obtained with optogenetics is more reliable.

A significant completion compared to our previous work was the introduction of CCK basket neurons. There are no direct data indicating the important role of these interneurons in the generation of theta rhythm, but there are many indirect ones. This group of neurons receives density input from the medial septum (Joshi et al., 2017). The discharges of this group of neurons occur in the ascending phase of the theta rhythm, i.e., during the period of hyperpolarization of the soma of pyramidal neurons (Klausberger et al., 2005; Somogyi et al., 2014), from which it is very logical to assume that they provide this hyperpolarization.

The literature presents opposite data on the role of NMDA receptors in theta rhythm. Some authors suggest a large contribution of NMDA receptors to theta rhythm, especially in the blockade of muscarinic receptors (Buzsáki, 2002; Soltesz & Deschênes, 1993). Another study has showed no effect of the NMDA receptor blocker on the theta rhythm power of the injected drug in the hippocampus (Gu, Alexander, Dudek, & Yakel, 2017). Based on this data, we have not introduce NMDA receptors, so as not to complicate the model.

To reproduce the experimental results in our model, the phase relations of the external inputs are extremely important. The phase difference between the inputs from the CA3 field and the MEC is known from experimental data (Mizuseki, Sirota, Pastalkova, & Buzsáki, 2009). It is also known that the inputs from the Teevra cells coupling to the minimum of the theta cycle (Joshi et al., 2017). The phase relations between the inputs from the CA3 field and the Teevra cells are very important for our model since they allow us to specify the excitation and inhibition of the perisomatic zone of pyramidal neurons. There is also a cholinergic input to OLM neurons via nicotine receptors (Haam, Zhou, Cui, & Yakel, 2018). Although there is currently no reliable data on the phase relationship of cholinergic neurons and theta rhythm, we assumed that it falls on the minimum of theta rhythm. This hypothesis allows us to strengthen the binding of OLM neurons to the minimum of the theta cycle, this input in our model has an auxiliary value.

### Key assumptions and predictions about the slow, middle, and fast gamma rhythms, ripple oscillations, and phase precession

Models of slow gamma rhythm, ripple oscillations, and phase precession are taken almost entirely from the cited works, so we can only refer to the originals to discuss these aspects. We will simply list the most important assumptions and consequences without a detailed discussion.

1. The pyramidal neurons that make up the active neural ensemble (currently active place cells) form a slow gamma rhythm by receiving more excitation from the CA3 field and the entorhinal cortex (Colgin et al., 2009; Fernández-Ruiz et al., 2017; Senior et al., 2008).
2. Spatial modulation of the PV basket neurons is formed by input from CA3 and local pyramidal neurons (Royer et al., 2012).
3. The phase precession is formed by the input from the entorhinal cortex and the CA3 field with a phase and time shift (Chance, 2012; Fernández-Ruiz et al., 2017; John O’Keefe & Burgess, 2005).

In our model, the middle gamma rhythm is formed by simple induction from the entorhinal cortex. The maximum power of the middle gamma rhythm is observed in the lacunosum molecular layer (Fernández-Ruiz et al., 2017). At the moment of arrival of the signal via the perforant path, a current sink is observed in the lacunosum molecular layer, which indicates the predominance of excitation over inhibition (Mizuseki et al., 2009). These two facts support a hypothesis that the middle gamma rhythm is formed by excitatory postsynaptic potentials from signals coming along the perforant path. In this process, it is possible to switch the signal from MEC via interneurons, but there is no data on this possibility. Since MEC neurons discharge at the ascending phase of the theta rhythm in the CA1 pyramidal layer, the most likely groups of neurons that discharge at this time and inhibit apical dendrites are ivy and neurogliaform interneurons. However, these inhibitory postsynaptic potentials on pyramidal neurons from the activation of these interneurons have very long characteristic times from 11 to 50 ms, which is significantly more than the period of the middle gamma rhythm (Krook-Magnuson, Luu, Lee, Varga, & Soltesz, 2011; Price et al., 2005).

The fast gamma rhythm in our model is provided by the action potentials of a large number of pyramidal neurons since the maximum excitation is observed at the minimum of the theta rhythm. Due to the overlap of excitatory inputs, a small group of pyramidal neurons gives synchronous action potentials, which is reflected in the field potential.

### Future research directions

We believe that the theoretical models proposed in neuroscience should follow the principle of Occam's razor. Models should aim to describe in the most detail the properties of an object or phenomenon observed experimentally, while theoretical models should be as simple as possible. To date, in theoretical neuroscience, simple models have been proposed for some relatively simple experimental phenomena. For example, about 6 theoretical models have been proposed to explain the effect of phase precession of place cells (Burgess & O’Keefe, 2011). We believe that the problem of constructing models of complex phenomena can be solved by sequentially combining simple models since a balance can be maintained between the complexity of described experimental phenomena and the complexity of a model. We have tried to do in this article to describe the rhythms in the CA1 field of the hippocampus. We believe that the generalization of the proposed model to the other regions of the hippocampal formation will be a promising continuation. The dentate gyrus, field CA3, and subiculum similar to the field CA1 by a set of populations of neurons, their connections, as well as the properties of the rhythms (Buzsáki, 2015; Carr et al., 2012; Mizuseki et al., 2009; White, Rees, Wheeler, Hamilton, & Ascoli, 2020). The model of generating rhythms in the neural network of the full hippocampus, taking into account the characteristics of different regions, will have great predictive and descriptive power.

Another promising direction for the development of the proposed ideas is to combine a rhythm model with models of place cells, time cells, and grid cells. As with rhythms, many models have been proposed to explain the individual effects associated with place cells (Burgess & O’Keefe, 2011). The problem with most of these models is their small relation to rhythmic processes, but there is a lot of experimental evidence of an indissoluble connection between cognitive processes and rhythms (Buzsáki & Moser, 2013; Buzsáki, 2015; Colgin, 2016; Vinogradova, 1995). Integrating a detailed model of rhythms with models that explain remapping, pattern separation, and pattern completion, and other correlates of cognitive functions at the neural level will be very important and fundamental.

## Acknowledgments

This work was supported by Grant 20-71-10109 of the Russian Science Foundation (RNF).

## Notes

### Competing Interest Statement

The authors have declared no competing interest.

