## supplemental materials for "A model of the CA1 field rhythms"

Mysin I.E.

#### 1 Programm instruments

We have used the simulator Neuron from Python. We have used the code of the models described in the article by Bezaire with coauthors [1] and in Cutsuridis and Poirazi [2]. Pyramidal and bistratified neurons were taken from the model of Cutsuridis and Poirazi, the source code of the models was taken from the repository `https://github.com/ModelDBRepository/181967`. The remaining neurons are taken from the model of Bezaire, the source code for the neuron models is taken from `https://github.com/ModelDBRepository/187604`. The source code of our model is available at `https://github.com/ivanmysin/CA1_rhythms_model`.

We have used libraries to process the simulated signals: NumPy [? ], Scipy [? ], Elephant [? ]. Graphs were plotted with Matplotlib [? ]. The code of processing is also available at `https://github.com/ivanmysin/CA1_rhythms_model`.

#### 2 Signal processing

We have processed the data using algorithms that are often used in experimental works. The filtration into frequency bands was carried out using a second-order Butterworth filter. The phase analysis of signals for finding the phase coupling of neurons was carried out using the Hilbert transform. Cross-rhythm phase-amplitude and phase-phase analysis has been performed in the same way as in the work (Belluscio et al. 2012) For phase-amplitude analysis, the phase of the theta rhythm was computed by filtering the signal in the theta range, Hilbert transform was followed by it. The power at gamma frequencies was found using the wavelet transform. The Morlet wavelet was used as the mother function, central frequency is 8 Hz. The phase-phase analysis of the coupling of theta and gamma rhythms was carried out by finding the signal phases in the theta and gamma ranges. Next, the phase of the signal from the theta band was multiplied by a series of coefficients, after which the phase difference was computed. After that, the phase array is averaged and the ray length is calculated. The modulation index was calculated using the Kullback-Leibler distance as described in (Tort et al. 2008). The amplitudes and phases for the modulation index were calculated using the wavelet transform. Phase precession analysis was performed using linear-circular linear regression (Kempter et al. 2012).

### 3 Models of artifitial spike generators

External inputs to the CA1 field are simulated with artificial spike generators. Almost all neurons in hippocampal formation are modulated by rhythms, therefore we have used von Misses distribution to take phase modulated spike trains:

$$f(\phi) = \frac{exp(\kappa \cdot cos(\phi - \mu))}{2\pi \cdot I_0(\kappa)} \tag{1}$$

where $I_0(\kappa)$ is the modified Bessel function of order 0. The parameters $\mu$ and $1/\kappa$ are analogous to the mean and variance in the normal distribution. Degree of neuron firing coupling to phase of thythm typicaly is described with parameter $R$ - ray length. Parameter $\kappa$ can be taken from $R$ by following equations [3]:

$$\kappa = \begin{cases} 2 \cdot R + R^3 + 5/6 \cdot R^5 & \text{if } R < 0.53 \\ -0.4 + 1.39 \cdot R + 0.43/(1 - R) & \text{if } 0.53 \leq R < 0.85 \\ 1/(3 \cdot R - 4 \cdot R^2 + R^3) & \text{if } R \geq 0.85 \end{cases} \tag{2}$$

A simple model of spike generator can be presented with an equation:

$$p_{sp} = \int_{\phi}^{\phi+\Delta\phi} f(\phi) d\phi \tag{3}$$

where $p_{sp}$ is the probabilty of spike generation during $\Delta\phi$. From phases we can go to the time with frequency ($\omega$):

$$\frac{d\phi}{dt} = 2\pi\omega, \ \phi(t) = 2\pi t\omega, \ d\phi = 2\pi\omega dt \tag{4}$$

we can join these equations:

$$p_{sp} = S \cdot \int_{t}^{t+\Delta t} f(2\pi t\omega) 2\pi\omega dt \tag{5}$$

$S$ is a normalization coefficient to get the given spike rate. Each generator is characterized by the preferred phase ($\mu$), frequency ($\omega$), ray length ($R = 0$ is mean that there is no coupling to phase, $R = 1$ is full coupling to phase $\mu$) and spike rate (regulated by $S$). We add also latency for the generator model, it is the period after spike generation with zero probability of new spike. We have used such models for inputs from the medial septum, the lateral entorhinal cortex, and the CA3 field.

Neurons of medial entorhinal cortex modulated by theta and gamma rhythms. We have simulated neurons of the MEC as grid cells. It means that they show slow periodic activity. It can be simulated as a product of probabilities simple oscillators:

$$p_{grid} = S \cdot p_{sp}(\omega_\theta, \mu_\theta, R_\theta) \cdot p_{sp}(\omega_\gamma, \mu_\gamma, R_\gamma) \cdot p_{sp}(\omega_{grid}, \mu_{grid}, R_{grid}) \qquad (6)$$

All parameters are given below. Input from the CA3 field is simulated with simple generators and with place cells. The CA3 neurons without place field activity are modulated only by theta rhythm. Place cells are modulated by theta and gamma rhythms, place field is simulated with gaussian:

$$p_{grid} = S \cdot p_{sp}(\omega_\theta, \mu_\theta, R_\theta) \cdot p_{sp}(\omega_\gamma, \mu_\gamma, R_\gamma) \cdot \frac{exp\left(\frac{-(t-t_{place})^2}{2 \cdot r_{place}^2}\right)}{2\pi r_{place}} \qquad (7)$$

where $t_{place}$ and $r_{place}$ are the center and the radius of a place field in time domain.

Table 1: Parameters of artifitial generators

| | ca3_non_spatial | msteevracells | msach |
|---|---|---|---|
| $R_\theta$ | 0.3 | 0.6 | 0.4 |
| $\omega_\theta,\ Hz$ | 7 | 7 | 7 |
| $latency,\ ms$ | 10 | 4 | 10 |
| $\mu_\theta,\ rad$ | 1.3 | $\pi$ | $\pi$ |
| $S$ | 1 | 15 | 3 |

Table 2: Parameters of artifitial generators

| | mec |
|---|---|
| $R_\gamma$ | 0.4 |
| $R_{grid}$ | 0.9 |
| $R_\theta$ | 0.3 |
| $\omega_{grid},\ Hz$ | 0.1 |
| $\omega_\gamma,\ Hz$ | 63 |
| $\mu_\gamma,\ rad$ | 0 |
| $latency,\ ms$ | 10 |
| $\omega_\theta,\ Hz$ | 7 |
| $\mu_\theta,\ rad$ | -1.04 |
| $S$ | 1e+08 |

Table 3: Parameters of artifitial generators

| | ca3_spatial |
|---|---|
| $R_\gamma$ | 0.4 |
| $R_\theta$ | 0.4 |
| $\omega_\gamma,\ Hz$ | 35 |
| $\mu_\gamma,\ rad$ | 0 |
| $latency,\ ms$ | 10 |
| $\omega_\theta,\ Hz$ | 7 |
| $\mu_\theta,\ rad$ | 1.3 |
| $r_{place},\ ms$ | 500 |
| $S$ | 1e+06 |

### 4 Neurons models

#### 4.1 General notes on neuron models

All neurons were multi compartments. The potential of each compartment is:

$$C\frac{dV}{dt} = -\sum I_{tm} - I_{syn} + I_{ext} \tag{8}$$

where $C$ is capacity in $\mu F/cm^2$, $V$ is potential in mV, $t$ is time in ms, $I$ is current in $\mu A/cm^2$. $I_{tm}$ are transmembrane currents, they are described in the following sections, and tables with parameters are given in 4.10. The common equation for transmembrane currents is:

$$I_{tm} = g_{max} \cdot x_1^N \cdot x_2^M \cdot x_3^K \cdot (V - E) \tag{9}$$

where $g_{max}$ is maximal coduntance in $mS/cm^2$, $E$ is reversal potential for this channel in $mV$. $x_1$, $x_2$, $x_3$ are the gate variables. All compartments have leak current:

$$I_L = g_L \cdot (V - E_L) \tag{10}$$

$g_L$ and $E_L$ are given in tables 4.10.

All gate variables were described with the classical equation:

$$\frac{dx}{dt} = \frac{x_\infty(V) - x}{\tau_x(V)} \tag{11}$$

The equations for $x_\infty(V)$ and $\tau_x(V)$ are given in the corresponding sections in explicit form or they can be given in form of functions $\alpha(V)$ and $\beta(V)$

$$\tau_x(V) = \frac{1}{\alpha(V) + \beta(V)} \quad x_\infty(V) = \alpha(V) \cdot \tau_x(V) \tag{12}$$

Numerical scheme:

$$x_{t+\Delta t} = x_t + \Big(1 - exp\Big(-\frac{\Delta t}{\tau_x}\Big)\Big) \cdot (x_\infty - x_t) \tag{13}$$

where $\Delta t = 0.1\ ms$ is integration step.

There is the function *Q(T)* in description of some currents, it is given by:

$$Q(T) = \frac{F}{R \cdot T} \tag{14}$$

where $R$ = *8.315 joule/deg*, $F$ = *9.648* $\cdot 10^4\ Coul$, $T$ is the temperature in degrees Kelvin. In all simulations *T=310 deg.*

$I_{syn}$ in equation (8) is a sum of synaptic current, which described in section 5.1.

External current ($I_{ext}$) in equation (8) simulates noise and influence of very long currents:

$$I_{ext} = I_{ext,mean} + \eta \cdot \sigma_{ext} \tag{15}$$

where $I_{ext,mean}$ is a constant value, $\eta$ is a random value from normal distribution with mean equals 0 and standard deviation equals 1, $\sigma_{ext} = 0.005\mu A/cm^2/\sqrt{ms}$ is std of noise. To simulate heterogeneously $I_{ext,mean}$ for each neuron is chosen from a lognormal distribution, a mean and std are given below.

#### 4.2 CA1 Pyramidal cell

Pyramid neurons were described using a 15-compartment model (figure 4). The model took into account soma, basal dendrites, and the proximal and distal parts of the apical dendrite.

Balance equation for the somatic compartment:

$$C\frac{dV_s}{dt} = -I_L - I_{Na} - I_{kdr} - I_A - I_M - I_H - I_{sAHP} - I_{mAHP} - I_{CaL} - \\ -I_{CaT} - I_{CaR} - I_{buff} - I_{syn} + I_{ext} \quad (16)$$

Balance equation for the axon:

$$C\frac{dV_a}{dt} = -I_L - I_{Na} - I_{kdr} - I_M - I_{syn} \quad (17)$$

Balance equation for the basal dendrites and the proximal part of the apical dendrite:

$$C\frac{dV_{rad,ori}}{dt} = -I_L - I_{Na} - I_{kdr} - I_A - I_M - I_H - I_{sAHP} - I_{mAHP} - \\ -I_{CaL} - I_{CaT} - I_{CaR} - I_{buff} - I_{syn} + I_{ext} \quad (18)$$

Balance equation for the distal part of the apical dendrite:

$$C\frac{dV_{LM}}{dt} = -I_L - I_{Na} - I_{kdr} - I_A - I_{syn} + I_{ext} \quad (19)$$

where $I_L$ is the leak current, $I_{Na}$ is the fast sodium current, $I_{kdr}$ is the delayed rectifier potassium current, $I_A$ is the A-type potassium current, $I_M$ is the M-typepotassium current, $I_H$ is a hyperpolarizing H-type current, $I_{CaL}$, $I_{CaT}$ and $I_{CaR}$ are the L-, T- and R-type $Ca^{2+}$ currents, respectively, $I_{sAHP}$ and $I_{mAHP}$ are slow and medium $Ca^{2+}$ activated $K^+$ currents, $I_{buff}$ is a calcium pump/buffering mechanism and $I_{syn}$ is the synaptic current. $I_{ext}$ is the tonic current and noise. The parameters for all ionic currents are listed in Table 4.

The sodium current is described by:

$$I_{Na} = g_{max,Na} \cdot m^2 \cdot h \cdot s \cdot (V - E_{Na}) \quad (20)$$

Activation and inactivation kinetics for $I_{Na}$ are given by:

$$m_\infty = \frac{1}{1 + exp(-\frac{V+40}{3})}, \quad \tau_m = 0.05\ ms \quad (21)$$

$$h_\infty = \frac{1}{1+exp(\frac{V+45}{3})},\ \tau_h = 0.5\ ms, \tag{22}$$

$$s_\infty = \frac{1+Na_{att}\cdot exp(\frac{V+60}{2})}{1+exp(\frac{V+60}{2})} \tag{23}$$

$$\tau_s = \frac{0.00333\cdot exp(0.0024\cdot(V+60)\cdot Q(T))}{1+exp(0.0012\cdot(V+60)\cdot Q(T))} \tag{24}$$

$Na_{att}$ variable represents the degree of sodium current attenuation and varies linearly from soma to distal trunk $Na_{att} \in [0, 1]$: 1 -maximum 0 - zero attenuation (Table 4). The delayed rectifier current is given by:

$$I_{Kdr} = g_{Kdr}\cdot m^2\cdot(V-E_K) \tag{25}$$

$$m_\infty = \frac{1}{1+exp(-\frac{V+42}{2})}, \tau_m = 2.2\ ms \tag{26}$$

The sodium and delayed rectifier channel properties are slightly different in the soma, axis and dendritic arbor. The $m_\infty$ and $h_\infty$ somatic/axonic HH channel kinetics as well as the time constants for both $I^{sa}_{Na}$ and $I^{sa}_{Kdr}$, are modified as follows. For the sodium:

$$m^{sa}_\infty = \frac{1}{1+exp(-\frac{V+44}{3})},\quad h^{sa}_\infty = \frac{1}{1+exp(\frac{V+49}{3.5})} \tag{27}$$

while for the potassium delayed rectifier

$$m^{sa}_\infty = \frac{1}{1+exp(-\frac{V+46.3}{3})} \tag{28}$$

The time constant for somatic and axonic $Na^+$ channel activation is kept the same $\tau_{\mathrm{m}} = 0.05$ ms while for inactivation is set to $\tau_{\mathrm{h}}= 1$ ms. The $\tau$-value for the delayed rectifier channel activation is set to $\tau_m = 3.5 ms$. The fast inactivating A-type $K^+$ current is described by

$$I_A = g_{max,A}\cdot n\cdot l\cdot(V-E_K) \tag{29}$$

$$n_\infty = \frac{\frac{-0.01(V+21.3)}{exp(-(V+21.3)/35)-1}}{\frac{-0.01(V+21.3)}{exp(-(V+21.3)/35)-1}+\frac{0.01(V+21.3)}{exp((V+21.3)/35)-1}}\quad \tau_n = 0.2\ ms \tag{30}$$

$$l_\infty = \frac{\frac{-0.01(V+58)}{exp((V+58)/8.2)-1}}{\frac{-0.01(V+58)}{exp((V+58)/8.2)-1}+\frac{0.01(V+58)}{exp(-(V+58)/8.2)-1}} \tag{31}$$

$$\tau_l = \begin{cases} 5 + 2.6(V + 20)/10 & \text{if } V > 20mV \\ 5 & \text{otherwise} \end{cases} \quad (32)$$

The hyperpolarizing H-current is:

$$I_H = g_{max,H} \cdot H \cdot (V - E_H) \quad (33)$$

$$H_\infty = \frac{1}{1 + exp((V - V_{half})/8)} \quad (34)$$

$$\tau_H = \frac{exp(0.003364 \cdot (V - V_{halft})}{0.05 \cdot q10^{(T-33)/10} \cdot (1 + a_H)} \quad (35)$$

$$a_H = exp(0.008316 \cdot (V - V_{halft})) \quad (36)$$

where $q10 = 0.4$, $V_{halft}$ see in Table 4.

The slowly activating voltage-dependent potassium current, $I_M$, is given by the equations:

$$I_m = 10^{-4} \cdot T_{adj}(T) \cdot g_m \cdot m \cdot (V - E_K) \quad (37)$$

$$\alpha_m = \frac{10^{-3} \cdot (V + 30)}{(1 - exp(-(V + 30)/9))T_{adj}(T)} \quad (38)$$

$$\beta_m = \frac{-10^{-3} \cdot (V + 30)}{(1 - exp((V + 30)/9))T_{adj}(T)} \quad (39)$$

$$T_{adj}(T) = 2.3^{(T-287)/10} \quad (40)$$

The slow after-hyperpolarizing current, is given by equations:

$$I_{sAHP} = g_{max,sAHP} \cdot m^3 \cdot (V - E_K) \quad (41)$$

$$\frac{dm}{dt} = \frac{\frac{Cac}{(1+Cac)} - m}{\tau_m} \quad (42)$$

$$\tau_m = max\Big(\frac{1}{0.003 \cdot (1 + Cac) \cdot 3^{(\deg C - 22)/10}},\ 0.5\Big) \quad (43)$$

where $Cac = (40 \cdot [Ca^{2+}]_{in})^2$.

The medium after-hyperpolarizing current, $\mathrm{I_{mAHP}}$ is:

$$I_{mAHP} = g_{max,mAHP} \cdot m \cdot (V - E_K) \quad (44)$$

$$\alpha_m(V) = \frac{0.48}{1 + \frac{0.18}{[Ca^{2+}]_{in}} \cdot exp(-1.68 \cdot V \cdot Q(T))} \tag{45}$$

$$\beta_m(V) = \frac{0.28}{1 + \frac{[Ca^{2+}]_{in}}{0.011 \cdot exp(-2 \cdot V \cdot Q(T))}} \tag{46}$$

Q(T) see formula (14).

The somatic high-voltage activated (HVA) L-type $Ca^{2+}$ current is given by

$$I^s_{CaL} = g^s_{max,CaL} \cdot m \cdot \frac{0.001 \cdot ghk(V, [Ca^{2+}]_{in}, [Ca^{2+}]_{out})}{0.001 + [Ca^{2+}]_{in}} \tag{47}$$

$$\alpha_m(V) = \frac{-0.275 \cdot (V + 27.01)}{exp(-(V + 27.01)/3.8) - 1} \tag{48}$$

$$\beta_m(V) = 4.7 \cdot exp(-(V + 63.01)/17) \tag{49}$$

whereas the dendritic L-type calcium channels have different kinetics:

$$I^d_{CaL} = g^d_{max,CaL} \cdot m^3 \cdot h \cdot (V - E_{Ca}) \tag{50}$$

$$m_\infty(V) = \frac{1}{1 + exp(-(V + 37))} \tag{51}$$

$$h_\infty(V) = \frac{1}{1 + exp((V + 41)/0.5)} \tag{52}$$

Their time constants are equal to $\tau_m$ = 3.6 $ms$ and $\tau_h$=29 $ms$. The low-voltage activated (LVA) T-type $Ca^{2+}$ channel kinetics are given by equations:

$$I_{CaT} = g_{max,CaT} \cdot m^2 \cdot h \cdot \frac{0.001 \cdot ghk(V, [Ca^{2+}]_{in}, [Ca^{2+}]_{out})}{0.001 + [Ca^{2+}]_{in}} \tag{53}$$

$$\begin{aligned} ghk(V,\ [Ca^{2+}]_{in},\ [Ca^{2+}]_{out}) = \\ = -x \cdot (1 - [Ca^{2+}]_{out}/[Ca^{2+}]_{in} \cdot exp(V/x)) \cdot f(V/x) \end{aligned} \tag{54}$$

$$x = \frac{0.0853 \cdot T}{2}, f(z) = \begin{cases} 1 - \frac{z}{2}, & if\ |z| < 10^{-4} \\ \frac{z}{e^z - 1}, & otherwise \end{cases} \tag{55}$$

$$\alpha_m(V) = -0.196 \cdot \frac{(V - 19.88)}{exp(-(V - 19.88)/10) - 1} \tag{56}$$

$$\beta_m(V) = 0.046 \cdot exp(-(V/22.73)) \tag{57}$$

$$\alpha_h(V) = 0.00011 \cdot exp(-(V + 57)/19) \tag{58}$$

$$\beta_h(V) = \frac{0.68}{exp(-(V - 15)/10) + 1} \tag{59}$$

where $[Ca^{2+}]_{in}$ and $[Ca^{2+}]_{out}$ are the internal and external calcium concentrations. The HVA R-type Ca2+ current is described by:

$$I_{CaR} = g_{max,CaR} \cdot m^3 \cdot h \cdot (V - E_{Ca}) \tag{60}$$

There is the difference between somatic and dendritic CaR currents in the $\alpha(V)$, $\beta(V)$ and $\tau$ parameter values. For the somatic current, $\tau_m = 100$ $ms$ and $\tau_h = 5ms$ while for the dendritic current $\tau_m = 50$ $ms$ and $\tau_h$=5 $ms$. The $\alpha(V)$ and $\beta(V)$ equations for dendritic CaR channels are:

$$m_\infty(V) = \frac{1}{1 + exp(-(V + 48.5)/3)} \tag{61}$$

$$h_\infty(V) = \frac{1}{1 + exp(V + 53)} \tag{62}$$

while for the CaR channels of the soma:

$$m_\infty(V) = \frac{1}{1 + exp(-(V + 60)/3)} \tag{63}$$

$$h_\infty(V) = \frac{1}{1 + exp(V + 62)} \tag{64}$$

There is a calcium pump/buffering mechanism in the cell body and along the apical and basal trunk. The factor for $Ca^{2+}$ entry was changed from $f_e =$ 10,000 to $f_e = 556$ and the rate of calcium removal was made 7 times faster. The kinetic equations are given by:

$$Ca_{inputflow} = \begin{cases} \frac{-f_e \cdot I_{Ca,sum}}{0.2 \cdot F}, & if\ Ca_{inputflow} > 0 \\ 0, & otherwise \end{cases} \tag{65}$$

$$\frac{d[Ca^{2+}]_{in}}{dt} = Ca_{inputflow} + \frac{[Ca^{2+}]_0 - [Ca^{2+}]_{in}}{\tau_{Ca}} \tag{66}$$

$[Ca^{2+}]_0 = 10^{-4}\ mM,\ \tau_{Ca} = 1400\ ms$

#### 4.3 Bistratified cells

Bistratified cells were described using a 13-compartment model. Balance equation for all compartments:

$$C\frac{dV}{dt} = -I_L - I_{Na} - I_{Kdr} - I_A - I_{CaL} - I_{CaN} - \\ -I_{AHP} - I_C - I_{syn} + I_{ext} \quad (67)$$

where $I_A$ is the A-type $K^+$ current,$I_{CaL}$ is the L-type $Ca^{2+}$ current, $I_{CaN}$ is the N-type $Ca^{2+}$ current, $I_{AHP}$ is the $Ca^{2+}$-dependent $K^+$ (SK) current, $I_C$ is the $Ca^{2+}$ and voltage-dependent $K^+$ (BK) current and $I_{syn}$ is the synaptic current. The conductances and reversal potentials of all ionic currents are listed in Table 12. The sodium current and its kinetics are described by:

$$I_{Na} = g_{max,Na} \cdot m^3 \cdot h \cdot (V - E_{Na}) \quad (68)$$

$$\alpha_m(V) = \frac{-0.3 \cdot (V - 25)}{1 - exp(-0.2 \cdot (V - 25))}, \quad \beta_m(V) = \frac{0.3 \cdot (V - 53)}{1 - exp(0.2 \cdot (V - 53))} \quad (69)$$

$$\alpha_h(V) = \frac{0.23}{exp((V - 3)/20)}, \quad \beta_h(V) = \frac{3.33}{1 + exp(-0.1 \cdot (V - 55.5))} \quad (70)$$

The fast delayed rectifier potassium current, $I_{Kdr}$ is given by:

$$I_{Kdr} = g_{max,Kdr} \cdot n^4 \cdot (V - E_K) \quad (71)$$

$$\alpha_n = \frac{-0.07 \cdot (V - 47)}{1 - exp((V - 47)/-6)}, \quad \beta_n = 0.264 \cdot exp((V - 22)/4) \quad (72)$$

The N-type calcium current, $I_{CaN}$, is given by

$$I_{CaN} = g_{max,CaN} \cdot c^2 \cdot d \cdot (V - E_{Ca}) \quad (73)$$

$$\alpha_c(V) = \frac{0.19 \cdot (19.88 - V)}{exp(0.1 \cdot (19.88 - V)) - 1}, \quad \beta_c(V) = 0.046 \cdot exp(-V/20.73) \quad (74)$$

$$\alpha_d = 1.6 \cdot 10^{-4} \cdot exp(-V/48.4), \quad \beta_d = \frac{1}{1 + exp(0.1 \cdot (39 - V))} \quad (75)$$

The $Ca^{2+}$-dependent $K^+$ (SK) current, $I_{AHP}$, is described by:

$$I_{AHP} = g_{max,AHP} \cdot q^2 \cdot (V - E_K) \tag{76}$$

$$\alpha_q(V) = \frac{0.00246}{exp((12 \cdot log_{10}([Ca^{2+}]_{in}) + 28.48)/-4.5)} \tag{77}$$

$$\beta_q(V) = \frac{0.006}{exp((12 \cdot log_{10}([Ca^{2+}]_{in}) + 60.4)/35)} \tag{78}$$

$$\frac{d[Ca^{2+}]_{in}}{dt} = B \sum_{T,N,L} I_{Ca} - \frac{[Ca^{2+}]_{in} - [Ca^{2+}]_0}{\tau_{Ca}} \tag{79}$$

where

$$B = 5.2 \cdot 10^{-6}/(A \cdot d) \tag{80}$$

in units of $mol/(Cm^3)$ for a shell of surface area $A$ and thickness $d$ (0.2 $\mu$m) and $\tau_{Ca}$ = 10 ms was the calcium removal rate. $[Ca^{2+}]_0$ = 5 $\mu$M was the resting calcium concentration. The $Ca^{2+}$ and voltage-dependent $K^+$ (BK) current, $I_c$, is:

$$I_C = g_{max,c} \cdot o \cdot (V - E_K) \tag{81}$$

$$\alpha_o = \frac{0.28 \cdot [Ca^{2+}]_{in}}{[Ca^{2+}]_{in} + 0.00048 \cdot exp(-1.68 \cdot F \cdot V/(R \cdot T))} \tag{82}$$

$$\beta_o = \frac{0.48}{1 + [Ca^{2+}]_{in}/(0.13 \cdot 10^{-6} \cdot exp(-2 \cdot F \cdot V/(R \cdot T)))} \tag{83}$$

The A-type $K^+$ current,$I_A$, is described by

$$I_A = g_{max,A} \cdot a \cdot b \cdot (V - E_K) \tag{84}$$

$$\alpha_a = \frac{0.02 \cdot (13.1 - V)}{exp(\frac{13.1-V}{10}) - 1},\ \beta_a = \frac{0.0175 \cdot (V - 40.1)}{exp(\frac{V-40.1}{10}) - 1} \tag{85}$$

$$\alpha_b = 0.0016 \cdot exp\Big(\frac{V+13}{-18}\Big),\ \beta_b = \frac{0.05}{1 + exp(\frac{10.1-V}{5})} \tag{86}$$

The L-type $Ca^{2+}$ current, $I_{CaL}$, is given by

$$I_{CaL} = g_{max,CaL} \cdot s_\infty^2 \cdot V \cdot \frac{1 - \frac{exp(2 \cdot F \cdot V/(k \cdot T)) \cdot [Ca^{2+}]_{in}}{[Ca^{2+}]_0}}{1 - exp(2 \cdot F \cdot V/(k \cdot T))} \tag{87}$$

where $F$ is Faraday's constant, $T$ is the temperature, $k$ is Boltzmann's constant, $[Ca^{2+}]_0$ is the equilibrium calcium concentration and $[Ca^{2+}]_{in}$ is described in equation (79). The activation variable, $s_\infty$, is then

$$\alpha_s(V) = \frac{15.69 \cdot (-V + 81.5)}{exp(\frac{-V+81.5}{10}) - 1}, \quad \beta_s(V) = 0.29 \cdot exp(-V/10.86) \tag{88}$$

#### 4.4 PV basket cells

PV basket cells were described with the 17-compartment model, 1 - soma, and 16 - dendrites (figure 1). Balance equation for all compartments:

$$C\frac{dV}{dt} = -I_L - I_{Na} - I_{Kdr} - I_{CaL} - I_{CaN} - I_{KCaB} - \\ -I_{KCaS} - I_{KA} - I_{syn} + I_{ext} \tag{89}$$

Sodium current for spike generation is:

$$I_{Na} = g_{max,Na} \cdot m^3 \cdot h \cdot (V - E_{Na}) \tag{90}$$

$$\alpha_m = \frac{-0.3 \cdot (V + 43)}{exp(-0.2 \cdot (V + 43)) - 1} \tag{91}$$

$$\beta_m = \frac{0.3 \cdot (V + 15)}{exp(0.2 \cdot (V + 15)) - 1} \tag{92}$$

$$\alpha_h = \frac{0.23}{exp(0.05 \cdot (V + 65))} \tag{93}$$

$$\beta_h = \frac{3.33}{exp(-0.1 \cdot (V + 12.5)) + 1} \tag{94}$$

Delayed rectification potassium current is

$$I_{Kdr} = g_{max,Kdr} \cdot n^4 \cdot (V - E_K) \tag{95}$$

$$\alpha_n = \frac{-0.07 \cdot (V + 18)}{exp(\frac{V+18}{-6}) - 1} \tag{96}$$

$$\beta_n = 0.264 \cdot exp\Big(\frac{V + 43}{40}\Big) \tag{97}$$

L-type of calcium current is:

$$I_{CaL} = g_{max,CaL} \cdot m^2 \cdot h \cdot ghk(V, [Ca_{2+}]_{in}, [Ca_{2+}]_{out}) \tag{98}$$

$$h = \frac{0.001}{0.001 + [Ca^{2+}]_{in}} \tag{99}$$

$$\alpha_m = \frac{15.69 \cdot (81.5 - V)}{exp(0.1(81.5 - V)) - 1} \tag{100}$$

$$\beta_m = 0.29 \cdot exp\Big(\frac{-V}{10.86}\Big) \tag{101}$$

$$ghk = -x \cdot \Big(1 - \Big(\frac{[Ca_{2+}]_{in}}{[Ca_{2+}]_{out}}\Big) \cdot exp\Big(\frac{V}{x}\Big)\Big) \frac{V}{x \cdot exp(V/x) - 1} \tag{102}$$

where

$$x = 0.04259 \cdot T \tag{103}$$

Calcium current N-type is given by following equation:

$$I_{CaN} = g_{max,CaN} \cdot c^2 \cdot d \cdot (V - E_{Ca}) \tag{104}$$

$$\alpha_c = \frac{-0.19 \cdot (V - 19.88)}{exp(0.1(V - 19.88)) - 1} \tag{105}$$

$$\beta_c = 0.046 \cdot exp\Big(\frac{-V}{20.73}\Big) \tag{106}$$

$$\alpha_d = 0.00016 \cdot exp\Big(\frac{-V}{48.4}\Big) \tag{107}$$

$$\beta_d = \frac{1}{exp(0.1 \cdot (39 - V)) + 1} \tag{108}$$

Calcium-dependent potassium current B:

$$I_{KCaB} = g_{max,KCaB} \cdot n \cdot (V - E_K) \tag{109}$$

$$\alpha_n = \frac{0.28 \cdot [Ca^{2+}]_{in}}{[Ca^{2+}]_{in} + 0.00048 \cdot exp(\frac{-1.68 \cdot F \cdot V}{R \cdot T})} \tag{110}$$

$$\beta_n = \frac{0.48}{1 + 130000 \cdot [Ca^{2+}]_{in} \cdot exp(\frac{2 \cdot F \cdot V}{R \cdot T})} \tag{111}$$

Calcium-dependent potassium current S:

$$I_{KCaS} = g_{max,KCaS} \cdot q^2 \cdot (V - E_K) \tag{112}$$

$$\alpha_q = q_{10} \cdot 15 \cdot ([Ca^{2+}]_{in})^2,\ \beta_q = q_{10} \cdot 0.00025 \tag{113}$$

$$q10 = 3^{0.1\cdot(T-307)} \tag{114}$$

A-type of potassium current is:

$$I_{KA} = g_{max,KA} \cdot n \cdot l \cdot (V - E_K) \tag{115}$$

$$n_\infty = \frac{1}{1 + exp(-21 \cdot (V + 33.6)/T)} \tag{116}$$

$$\tau_n = \frac{exp(-21 \cdot (V + 33.6)/T)}{q_{10} \cdot 0.02 \cdot (1 + exp(-21 \cdot (V + 33.6)/T))} \tag{117}$$

$$l_\infty = \frac{1}{1 + exp(46.41 \cdot (V + 83)/T)} \tag{118}$$

$$\tau_l = \frac{exp(46.41 \cdot (V + 83)/T)}{q_{10} \cdot 0.08 \cdot (1 + exp(46.41 \cdot (V + 83)/T))} \tag{119}$$

$$q10 = 3^{0.1\cdot(T-303)} \tag{120}$$

#### 4.5 CCK basket cells

CCK basket neurons as well as parvalbumin-containing ones were described by a model of 17 compartments, 1-soma, and 16 compartments of dendrites (figure 1). The potential of all compartments was described by the following equation:

$$\begin{aligned} C\frac{dV}{dt} = &-I_L - I_{Na} - I_{Kdr} - I_{CaL} - I_{CaN} - I_H - I_{KCaS} - \\ &-I_{KA} - I_{KCaB} - I_{KGroup} - I_{syn} + I_{ext} \end{aligned} \tag{121}$$

Sodium current is:

$$I_{Na} = g_{max,Na} \cdot m^3 \cdot h \cdot s \cdot (V - E_{Na}) \tag{122}$$

$$\alpha_m = \frac{-0.5 \cdot (V + 42)}{exp(-0.2 \cdot (V + 42)) - 1} \tag{123}$$

$$\beta_m = \frac{0.3 \cdot (V + 13)}{exp(0.2 \cdot (V + 13)) - 1} \tag{124}$$

$$\alpha_h = \frac{0.6}{exp(0.05 \cdot (V + 65))} \tag{125}$$

$$\beta_h = \frac{1.31}{exp(-0.1 \cdot (V + 12.5)) + 1} \tag{126}$$

$$\alpha_s = \frac{0.003}{exp(\frac{V+45}{6})} \tag{127}$$

$$\beta_s = \frac{0.005}{exp(-0.05 \cdot (V + 35))} \tag{128}$$

H-current is:

$$I_H = g_{max,H} \cdot H^2 \cdot (V - E_H) \tag{129}$$

$$\tau_H = \frac{1}{q_{10}} \cdot \left(120 + \frac{129.5}{1 + exp(1.2\ (V + 59.3))}\right) \tag{130}$$

$$H_\infty = \frac{1}{1 + exp(0.1\ (V + 91))} \tag{131}$$

$$q10 = 3^{0.1 \cdot (T - 307)} \tag{132}$$

Group potassium current:

$$I_{KGroup} = g_{max,KGroup} \cdot n \cdot (V - E_K) \tag{133}$$

$$\alpha_n(V) = \frac{-0.0189324 \cdot (V - 4.18371)}{exp(-0.15562 \cdot (V - 4.18371)) - 1} \tag{134}$$

$$\beta_n(V) = 0.015857 \cdot exp\left(\frac{-V}{25.4834}\right) \tag{135}$$

$I_{Kdr}, I_{CaL}, I_{CaN}, I_{KCaS}, I_{KCaB}, I_{KA}$ were same as PV basket cells and described equations (95, 98, 104, 112, 109, 115) respectivly.

#### 4.6 OLM cells

OLM neurons were modeled using 4 compartments - 1 soma, 1 axon, and 2 dendritic compartments (figure 2). Potential equation for the neuron soma:

$$C\frac{dV_s}{dt} = -I_L - I_{Na} - I_{Kdr} - I_H - I_{KA} - \\ -I_{syn} + I_{ext} \tag{136}$$

The equation for the dendrites:

$$C\frac{dV_d}{dt} = -I_L - I_{Na} - I_{Kdr} - I_{KA} - \\ -I_{syn} + I_{ext} \tag{137}$$

The equation for the axon:

$$C\frac{dV_a}{dt} = -I_L - I_{Na} - I_{Kdr} - I_{syn} \tag{138}$$

Sodium currents and potassium delayed rectification currents were modeled in the same way as in PV basket neurons, see equations (90) and (95) respectively.

H-current is:

$$I_H = g_{max,H} \cdot H \cdot (V - E_H) \tag{139}$$

$$H_\infty = \frac{1}{1 + exp(0.98 \cdot (V + 84.1))} \tag{140}$$

$$\tau_H = 100 + \frac{1}{exp(-(17.9 + 0.116 \cdot V)) + exp(0.09 \cdot V - 1.84)} \tag{141}$$

Potassium current A-type is:

$$I_{KA} = g_{max,KA} \cdot a \cdot b \cdot (V - E_K) \tag{142}$$

$$a_\infty = \frac{1}{1 + exp(\frac{V+14}{-16.6})} \quad \tau_a = 5 \tag{143}$$

$$b_\infty = \frac{1}{1 + exp(\frac{V+71}{7.3})} \tag{144}$$

$$\tau_b = \frac{1}{\frac{0.000009}{exp(\frac{V-26}{18.5})} + \frac{0.014}{exp(\frac{V+70}{-11})+0.2}} \tag{145}$$

#### 4.7 Axo-axonic cells

Axo-axonal interneurons were modeled using a 17-compartment model: 1 soma and 16 compartments for dendrites (figure 1). All compartments were described by the same set of channels:

$$C\frac{dV}{dt} = -I_L - I_{Na} - I_{Kdr} - I_{CaL} - I_{CaN} - \\ -I_{KCaS} - I_{KA} - I_{KCaB} - I_{syn} + I_{ext} \tag{146}$$

The description of all currents is similar to those used above for other neurons, see equations (90, 95, 98, 104, 112, 115)

#### 4.8 Ivy and neurogiaform cells

Ivy and neurogliaformes were described using 17 compartments: 1 soma and 16 dendritic compartments (figure 1). All the compartments in both types of neurons had the same set of channels:

$$C\frac{dV}{dt} = -I_L - I_{Na} - I_{Kdr} - I_{CaL} - I_{CaN} - \\ -I_{KCaS} - I_{KA} - I_{KCaB} - I_{syn} + I_{ext} \tag{147}$$

Sodium channels:

$$I_{Na} = g_{max,Na} \cdot m^3 \cdot h \cdot (V - E_{Na}) \tag{148}$$

$$\alpha_m = \frac{-0.34133 \cdot (V + 24)}{exp(-0.2 \cdot (V + 24)) - 1} \tag{149}$$

$$\beta_m = \frac{0.28483 \cdot (V - 4)}{exp(0.2 \cdot (V - 4)) - 1} \tag{150}$$

$$\alpha_h = \frac{0.29648}{exp(0.05 \cdot (V + 64.4184))} \tag{151}$$

$$\beta_h = \frac{3.0931}{exp(-0.1 \cdot (V + 12.1463)) + 1} \tag{152}$$

$I_{Kdr}$:

$$I_{Kdr} = g_{max,Kdr} \cdot n^4 \cdot (V - E_K) \tag{153}$$

$$\alpha_n = \frac{-0.07(V + 8)}{exp(\frac{V+8}{-6}) - 1} \tag{154}$$

$$\beta_n = 0.264 \cdot exp\Big(\frac{V + 33}{40}\Big) \tag{155}$$

A-type of potassium current:

$$I_{KA} = g_{max,KA} \cdot n \cdot l \cdot (V - E_K) \tag{156}$$

$$n_\infty = \frac{1}{1 + exp(-34.8 \cdot (V + 23.6)/T)} \tag{157}$$

$$\tau_n = \frac{exp(-34.8 \cdot (V + 23.6))}{0.02 \cdot q_{10} \cdot (1 + exp(-34.8 \cdot (V + 23.6)/T))} \tag{158}$$

$$l_\infty = \frac{1}{1 + exp \cdot (46.41 \cdot (V + 83)/T)} \tag{159}$$

$$\tau_l = \frac{exp(46.41 \cdot (V+83))}{0.08 \cdot q_{10} \cdot (1 + exp(46.41 \cdot (V+83)/T))} \tag{160}$$

$$q10 = 3^{0.1 \cdot (T-303)} \tag{161}$$

The description of all other currents is similar to those used above for other neurons, see equations (98, 104, 112, 109)

#### 4.9 Schaffer collateral associated cells

SCA cells were described using 17 compartments: 1 soma and 16 dendritic compartments (figure 1). All the compartments were described using the following balance equation:

$$C\frac{dV_s}{dt} = -I_L - I_{Na} - I_{Kdr} - I_{CaL} - I_{CaN} - I_H - I_{KCaS} - \\ -I_{KA} - I_{KCaB} - I_{KGroup} - I_{syn} + I_{ext} \tag{162}$$

Sodium current, $I_{KGroup}$ and $I_H$ are described by same way as for CCK basket cells - equation (122, 133, 129), equations for another currents are identical equations for PV basket cells (98, 104, 112, 115)

#### 4.10 Tables of neuronal parameters

Table 4: Pyramidal cells parameters

| Parameters | Soma | Axon | OriProx | OriDist | RadProx | RadMed | RadDist | LM |
|---|---|---|---|---|---|---|---|---|
| $C,\ \mu F/cm2$ | 1 | 1 | 1 | 1 | 1 | 1 | 1 | 1 |
| $Ra,\ Ohm \cdot cm$ | 50 | 50 | 50 | 50 | 50 | 50 | 50 | 50 |
| $g_L, mmS/cm2$ | 0.2 | 0.005 | 0.005 | 0.005 | 0.005 | 0.005 | 0.005 | 0.005 |
| $g_{max,Na} mS/cm2$ | 7 | 100 | 7 | 7 | 7 | 7 | 7 | 7 |
| $g_{max,Kdr}, mS/cm2$ | 1.4 | 20 | 0.868 | 0.868 | 0.868 | 0.868 | 0.868 | 0.868 |
| $g_{max,KA}, mS/cm2$ in proximal dendrites | 2.5 | — | 7.5 | 7.5 | 0.015 | — | — | — |
| $g_{max,KA}, mS/cm2$ in distal dendrites | — | — | — | — | — | 30 | 45 | 49 |
| $g_{max,KM}, mS/cm2$ | 60 | 30 | 60 | 60 | 60 | 60 | 60 | — |
| $g_{max,H}, mS/cm2$ | 0.05 | — | 0.05 | 0.1 | 0.1 | 0.2 | 0.35 | — |
| $V_{half}, mV$ | -73 | — | -81 | -81 | -82 | -81 | -81 | — |
| $g_{max,CaL}, mS/cm2$ | 0.7 | — | 0.031635 | 0.031635 | 0.031635 | 3.1635 | 3.1635 | — |

Continued on next page

Table 4: Pyramidal cells parameters

| Parameters | Soma | Axon | OriProx | OriDist | RadProx | RadMed | RadDist | LM |
|---|---|---|---|---|---|---|---|---|
| $g_{max,CaT}, mS/cm2$ | 0.05 | — | 0.1 | 0.1 | 0.1 | 0.1 | 0.1 | — |
| $g_{max,CaR}, mS/cm2$ | 0.3 | — | 0.03 | 0.03 | 0.03 | 0.03 | 0.03 | — |
| $g_{max,sAHP}, mS/cm2$ | 0.0005 | — | 0.5 | 0.5 | 0.5 | 0.5 | 0.5 | — |
| $g_{max,mAHP}, mS/cm2$ | 0.09075 | — | 33 | 33 | 33 | 33 | 4.1 | — |
| $E_L,\ mV$ | -70 | -70 | -70 | -70 | -70 | -70 | -70 | -70 |
| $E_{Na},\ mV$ | 50 | 50 | 50 | 50 | 50 | 50 | 50 | 50 |
| $E_H,\ mV$ | -10 | — | -10 | -10 | -10 | -10 | -10 | -10 |
| $E_{Ca},\ mV$ | 140 | — | 140 | 140 | 140 | 140 | 140 | — |
| $E_K,\ mV$ | -80 | -80 | -80 | -80 | -80 | -80 | -80 | -80 |

Table 5: PV basket cells parameters

| Parameters | soma | dend 1 | dend 2 | dend 3 | dend 4 | dend 5 | dend 6 | dend 7 | dend 8 |
|---|---|---|---|---|---|---|---|---|---|
| $L,\ \mu m$ | 20 | 100 | 100 | 200 | 100 | 100 | 100 | 100 | 100 |
| $Ra, ohm \cdot cm$ | 100 | 100 | 100 | 100 | 100 | 100 | 100 | 100 | 100 |
| $C, \mu F/cm^2$ | 1.4 | 1.4 | 1.4 | 1.4 | 1.4 | 1.4 | 1.4 | 1.4 | 1.4 |
| $D, \mu m$ | 10 | 4 | 3 | 2 | 1.5 | 1 | 2 | 1.5 | 1 |
| $E_L, mV$ | -65 | -65 | -65 | -65 | -65 | -65 | -65 | -65 | -65 |
| $g_{max,CaL}\ mS/cm^2$ | 5 | 5 | 5 | 5 | 5 | 5 | 5 | 5 | 5 |
| $g_{max,CaN}\ mS/cm^2$ | 0.8 | 0.8 | 0.8 | 0.8 | 0.8 | 0.8 | 0.8 | 0.8 | 0.8 |
| $g_{max,KCaS}\ mS/cm^2$ | 0.002 | 0.002 | 0.002 | 0.002 | 0.002 | 0.002 | 0.002 | 0.002 | 0.002 |
| $g_{max,Kdr}\ mS/cm^2$ | 13 | 13 | 13 | 13 | 13 | 13 | 13 | 13 | 13 |
| $g_{max,KA}\ mS/cm^2$ | 0.15 | 0.15 | 0.15 | 0.15 | 0.15 | 0.15 | 0.15 | 0.15 | 0.15 |
| $g_{max,KCaB}\ mS/cm^2$ | 0.0002 | 0.0002 | 0.0002 | 0.0002 | 0.0002 | 0.0002 | 0.0002 | 0.0002 | 0.0002 |
| $g_{max,Na}\ mS/cm^2$ | 150 | 150 | 150 | 150 | 150 | 150 | 150 | 150 | 150 |
| $g_L, mS/cm^2$ | 0.18 | 0.18 | 0.18 | 0.18 | 0.18 | 0.18 | 0.18 | 0.18 | 0.18 |

Table 6: Axo-axonal cells parameters

| Parameters | soma | dend 1 | dend 2 | dend 3 | dend 4 | dend 5 | dend 6 | dend 7 | dend 8 |
|---|---|---|---|---|---|---|---|---|---|
| $L,\ \mu m$ | 20 | 100 | 100 | 200 | 100 | 100 | 100 | 100 | 100 |
| $Ra, ohm \cdot cm$ | 100 | 100 | 100 | 100 | 100 | 100 | 100 | 100 | 100 |
| $C, \mu F/cm^2$ | 1.4 | 1.4 | 1.4 | 1.4 | 1.4 | 1.4 | 1.4 | 1.4 | 1.4 |
| $D, \mu m$ | 10 | 4 | 3 | 2 | 1.5 | 1 | 2 | 1.5 | 1 |
| $E_L, mV$ | -65 | -65 | -65 | -65 | -65 | -65 | -65 | -65 | -65 |
| $g_{max,CaL}\ mS/cm^2$ | 5 | 5 | 5 | 5 | 5 | 5 | 5 | 5 | 5 |
| $g_{max,CaN}\ mS/cm^2$ | 0.8 | 0.8 | 0.8 | 0.8 | 0.8 | 0.8 | 0.8 | 0.8 | 0.8 |
| $g_{max,KCaS}\ mS/cm^2$ | 0.002 | 0.002 | 0.002 | 0.002 | 0.002 | 0.002 | 0.002 | 0.002 | 0.002 |

Continued on next page

Table 6: Axo-axonal cells parameters

| Parameters | soma | dend 1 | dend 2 | dend 3 | dend 4 | dend 5 | dend 6 | dend 7 | dend 8 |
|---|---|---|---|---|---|---|---|---|---|
| $g_{max,Kdr}$ $mS/cm^2$ | 13 | 13 | 13 | 13 | 13 | 13 | 13 | 13 | 13 |
| $g_{max,KA}$ $mS/cm^2$ | 0.15 | 0.15 | 0.15 | 0.15 | 0.15 | 0.15 | 0.15 | 0.15 | 0.15 |
| $g_{max,KCaB}$ $mS/cm^2$ | 0.0002 | 0.0002 | 0.0002 | 0.0002 | 0.0002 | 0.0002 | 0.0002 | 0.0002 | 0.0002 |
| $g_{max,Na}$ $mS/cm^2$ | 150 | 150 | 150 | 150 | 150 | 150 | 150 | 150 | 150 |
| $g_L, mS/cm^2$ | 0.18 | 0.18 | 0.18 | 0.18 | 0.18 | 0.18 | 0.18 | 0.18 | 0.18 |

Table 7: CCK basket cells parameters

| Parameters | soma | dend 1 | dend 2 | dend 3 | dend 4 | dend 5 | dend 6 | dend 7 | dend 8 |
|---|---|---|---|---|---|---|---|---|---|
| $L,\ \mu m$ | 20 | 100 | 100 | 200 | 100 | 100 | 100 | 100 | 100 |
| $Ra, ohm \cdot cm$ | 150 | 150 | 150 | 150 | 150 | 150 | 150 | 150 | 150 |
| $C, \mu F/cm^2$ | 1.4 | 1.4 | 1.4 | 1.4 | 1.4 | 1.4 | 1.4 | 1.4 | 1.4 |
| $D, \mu m$ | 10 | 3.5 | 2.5 | 1.5 | 1.2 | 1 | 1.5 | 1.2 | 1 |
| $E_H, mV$ | 0 | 0 | 0 | 0 | 0 | 0 | 0 | 0 | 0 |
| $E_L, mV$ | -72 | -72 | -72 | -72 | -72 | -72 | -72 | -72 | -72 |
| $g_{max,CaL}$ $mS/cm^2$ | 2.7 | 2.7 | 2.7 | 2.7 | 2.7 | 2.7 | 2.7 | 2.7 | 2.7 |
| $g_{max,CaN}$ $mS/cm^2$ | 0.02 | 0.02 | 0.02 | 0.02 | 0.02 | 0.02 | 0.02 | 0.02 | 0.02 |
| $g_{max,H}$ $mS/cm^2$ | 0.1 | 0.1 | 0.1 | 0.1 | 0.1 | 0.1 | 0.1 | 0.1 | 0.1 |
| $g_{max,KCaS}$ $mS/cm^2$ | 0.004 | 0.004 | 0.004 | 0.004 | 0.004 | 0.004 | 0.004 | 0.004 | 0.004 |
| $g_{max,Kdr}$ $mS/cm^2$ | 0.008 | 0.08 | — | — | — | — | 0.08 | — | — |
| $g_{max,KA}$ $mS/cm^2$ | 0.4 | 0.4 | 0.4 | 0.4 | 0.4 | 0.4 | 0.4 | 0.4 | 0.4 |
| $g_{max,KCaB}$ $mS/cm^2$ | 0.04 | 0.04 | 0.04 | 0.04 | 0.04 | 0.04 | 0.04 | 0.04 | 0.04 |
| $g_{max,KGroup}$ $mS/cm^2$ | 1.3 | 2.6 | — | — | — | — | 2.6 | — | — |
| $g_{max,Na}$ $mS/cm^2$ | 18 | 9 | — | — | — | — | 9 | — | — |
| $g_L, mS/cm^2$ | 0.037 | 0.037 | 0.037 | 0.037 | 0.037 | 0.037 | 0.037 | 0.037 | 0.037 |

Table 8: OLM cells parameters

| Parameters | soma | dend 1 | dend 2 | axon |
|---|---|---|---|---|
| $L,\ \mu m$ | 20 | 250 | 250 | 150 |
| $Ra, ohm \cdot cm$ | 150 | 150 | 150 | 150 |
| $C, \mu F/cm^2$ | 1.3 | 1.3 | 1.3 | 1.3 |
| $D, \mu m$ | 10 | 3 | 3 | 1.5 |
| $E_L, mV$ | -67 | -67 | -67 | -67 |
| $g_{max,H}$ $mS/cm^2$ | 0.5 | — | — | — |

Continued on next page

Table 8: OLM cells parameters

| Parameters | soma | dend 1 | dend 2 | axon |
|---|---|---|---|---|
| $g_{max,Kdr}$ $mS/cm^2$ | 73 | 110 | 110 | 120 |
| $g_{max,KA}$ $mS/cm^2$ | 5 | 2.8 | 2.8 | — |
| $g_{max,Na}$ $mS/cm^2$ | 11 | 23 | 23 | 17 |
| $g_L, mS/cm^2$ | 0.01 | 0.01 | 0.01 | 0.01 |

Table 9: Neurogialform cells parameters

| Parameters | soma | dend 1 | dend 2 | dend 3 | dend 4 | dend 5 | dend 6 | dend 7 | dend 8 |
|---|---|---|---|---|---|---|---|---|---|
| L | 20 | 100 | 100 | 200 | 100 | 100 | 100 | 100 | 100 |
| $Ra, ohm \cdot cm$ | 14 | 14 | 14 | 14 | 14 | 14 | 14 | 14 | 14 |
| $C, \mu F/cm^2$ | 1.8 | 1.8 | 1.8 | 1.8 | 1.8 | 1.8 | 1.8 | 1.8 | 1.8 |
| $D, \mu m$ | 10 | 4 | 3 | 2 | 1.5 | 1 | 2 | 1.5 | 1 |
| $E_L, mV$ | -60 | -60 | -60 | -60 | -60 | -60 | -60 | -60 | -60 |
| $g_{max,CaL}$ $mS/cm^2$ | 56 | 56 | 56 | 56 | 56 | 56 | 56 | 56 | 56 |
| $g_{max,CaN}$ $mS/cm^2$ | 0.58 | 0.58 | 0.58 | 0.58 | 0.58 | 0.58 | 0.58 | 0.58 | 0.58 |
| $g_{max,KCaS}$ $mS/cm^2$ | 0.00045 | 0.00045 | 0.00045 | 0.00045 | 0.00045 | 0.00045 | 0.00045 | 0.00045 | 0.00045 |
| $g_{max,Kdr}$ $mS/cm^2$ | 160 | 2.2 | — | — | — | — | 2.2 | — | — |
| $g_{max,KA}$ $mS/cm^2$ | 0.0052 | 0.0052 | 0.0052 | 0.0052 | 0.0052 | 0.0052 | 0.0052 | 0.0052 | 0.0052 |
| $g_{max,KCaB}$ $mS/cm^2$ | 0.001 | 0.001 | 0.001 | 0.001 | 0.001 | 0.001 | 0.001 | 0.001 | 0.001 |
| $g_{max,Na}$ $mS/cm^2$ | 3800 | 260 | — | — | — | — | 260 | — | — |
| $g_L, mS/cm^2$ | 0.085 | 0.085 | 0.085 | 0.085 | 0.085 | 0.085 | 0.085 | 0.085 | 0.085 |

Table 10: Ivy cells parameters

| Parameters | soma | dend 1 | dend 2 | dend 3 | dend 4 | dend 5 | dend 6 | dend 7 | dend 8 |
|---|---|---|---|---|---|---|---|---|---|
| L | 20 | 100 | 100 | 200 | 100 | 100 | 100 | 100 | 100 |
| $Ra, ohm \cdot cm$ | 14 | 14 | 14 | 14 | 14 | 14 | 14 | 14 | 14 |
| $C, \mu F/cm^2$ | 1.8 | 1.8 | 1.8 | 1.8 | 1.8 | 1.8 | 1.8 | 1.8 | 1.8 |
| $D, \mu m$ | 10 | 4 | 3 | 2 | 1.5 | 1 | 2 | 1.5 | 1 |
| $E_L, mV$ | -60 | -60 | -60 | -60 | -60 | -60 | -60 | -60 | -60 |
| $g_{max,CaL}$ $mS/cm^2$ | 56 | 56 | 56 | 56 | 56 | 56 | 56 | 56 | 56 |
| $g_{max,CaN}$ $mS/cm^2$ | 0.58 | 0.58 | 0.58 | 0.58 | 0.58 | 0.58 | 0.58 | 0.58 | 0.58 |
| $g_{max,KCaS}$ $mS/cm^2$ | 0.00045 | 0.00045 | 0.00045 | 0.00045 | 0.00045 | 0.00045 | 0.00045 | 0.00045 | 0.00045 |
| $g_{max,Kdr}$ $mS/cm^2$ | 160 | 2.2 | — | — | — | — | 2.2 | — | — |
| $g_{max,KA}$ $mS/cm^2$ | 0.0052 | 0.0052 | 0.0052 | 0.0052 | 0.0052 | 0.0052 | 0.0052 | 0.0052 | 0.0052 |
| $g_{max,KCaB}$ $mS/cm^2$ | 0.001 | 0.001 | 0.001 | 0.001 | 0.001 | 0.001 | 0.001 | 0.001 | 0.001 |
| $g_{max,Na}$ $mS/cm^2$ | 3800 | 260 | — | — | — | — | 260 | — | — |
| $g_L, mS/cm^2$ | 0.085 | 0.085 | 0.085 | 0.085 | 0.085 | 0.085 | 0.085 | 0.085 | 0.085 |

Table 11: Schaffer collateral-associated cells parameters

| Parameters | soma | dend 1 | dend 2 | dend 3 | dend 4 | dend 5 | dend 6 | dend 7 | dend 8 |
|---|---|---|---|---|---|---|---|---|---|
| $L,\ \mu m$ | 10 | 100 | 100 | 200 | 100 | 100 | 100 | 100 | 100 |
| $Ra, ohm \cdot cm$ | 150 | 150 | 150 | 150 | 150 | 150 | 150 | 150 | 150 |
| $C, \mu F/cm^2$ | 1.2 | 1.2 | 1.2 | 1.2 | 1.2 | 1.2 | 1.2 | 1.2 | 1.2 |
| $D, \mu m$ | 10 | 3.5 | 2.5 | 1.5 | 1.2 | 1 | 1.5 | 1.2 | 1 |
| $E_H, mV$ | 0 | 0 | 0 | 0 | 0 | 0 | 0 | 0 | 0 |
| $E_L, mV$ | -72 | -72 | -72 | -72 | -72 | -72 | -72 | -72 | -72 |
| $g_{max,CaL}\ mS/cm^2$ | 1 | 1 | 1 | 1 | 1 | 1 | 1 | 1 | 1 |
| $g_{max,CaN}\ mS/cm^2$ | 0.02 | 0.02 | 0.02 | 0.02 | 0.02 | 0.02 | 0.02 | 0.02 | 0.02 |
| $g_{max,H}\ mS/cm^2$ | 0.07 | 0.07 | 0.07 | 0.07 | 0.07 | 0.07 | 0.07 | 0.07 | 0.07 |
| $g_{max,KCaS}\ mS/cm^2$ | 0.001 | 0.001 | 0.001 | 0.001 | 0.001 | 0.001 | 0.001 | 0.001 | 0.001 |
| $g_{max,Kdr}\ mS/cm^2$ | 0.006 | 0.06 | — | — | — | — | 0.06 | — | — |
| $g_{max,KA}\ mS/cm^2$ | 0.1 | 0.1 | 0.1 | 0.1 | 0.1 | 0.1 | 0.1 | 0.1 | 0.1 |
| $g_{max,KCaB}\ mS/cm^2$ | 0.007 | 0.007 | 0.007 | 0.007 | 0.007 | 0.007 | 0.007 | 0.007 | 0.007 |
| $g_{max,KGroup}\ mS/cm^2$ | 1.1 | 2.2 | — | — | — | — | 2.2 | — | — |
| $g_{max,Na}\ mS/cm^2$ | 40 | 20 | — | — | — | — | 20 | — | — |
| $g_L, mS/cm^2$ | 0.029 | 0.029 | 0.029 | 0.029 | 0.029 | 0.029 | 0.029 | 0.029 | 0.029 |

Table 12: Bistratifired cells parameters

| Parameters | All compartments |
|---|---|
| $C,\ \mu F/cm2$ | 1.4 |
| $Ra,\ Ohm \cdot cm$ | 100 |
| $g_{max,L},\ mS/cm2$ | 0.18 |
| $g_{max,Na},\ mS/cm2$ | 300 |
| $g_{max,Kdr},\ mS/cm2$ | 13 |
| $g_{max,KA},\ mS/cm2$ | 0.15 |
| $g_{max,CaL},\ mS/cm2$ | 5 |
| $g_{max,CaN},\ mS/cm2$ | 0.8 |
| $g_{max,AHP},\ mS/cm2$ | 0.002 |
| $g_{max,C},\ mS/cm2$ | 0.2 |
| $\tau_{Ca},\ ms$ | 10 |
| $E_{Na},\ mV$ | 55 |
| $E_K,\ mV$ | -90 |
| $E_{Ca},\ mV$ | 130 |
| $E_L,\ mV$ | -60 |
| $[Ca^{2+}]_0,\ mM$ | 0.005 |

### 5 Synapse models

#### 5.1 General description of synapse models

We simulate connections via gap junctions and chemical synapses with AMPAR, nicotinic and GABA-A receptors. The model of gap junctions is described in section (6). Current through all synaptic channels is simulated as:

$$I_{syn} = g_{max,syn} \cdot g \cdot (V - E_{syn}) \tag{163}$$

where $g_{max,syn}$ is maximal synaptic conductance $mS/cm^2$, $g$ is analog of gate variable for synaptic current. $E_{syn}$ is reversal potential in $mV$, for excitatory synapses with AMPA and nicotinic receptors $E_{syn} = 0\, mV$, for GABA-A channels $E_{GABA} = -75\, mV$. Dynamic of $g$ is simulated with the double exponential model:

$$g = exp\Big(\frac{t - t_0}{\tau_{rise}}\Big) - exp\Big(\frac{t - t_0}{\tau_{decay}}\Big) \tag{164}$$

where $t_0$ is a time of spike on presynaptic neuron with delay.

Parameters $\tau_{rise}$, $\tau_{decay}$ for synapses between each group of neurons are described in the following sections. For each synapse $g_{max,syn}$ and delay are chosen from a lognormal distribution, mean and std are given below.

#### 5.2 Synapses from pyramidal cells

Table 13: Synaptic connections to aac

| | $g_{max,mean}$ | $g_{max,\sigma}$ | $\tau_{rise}$ | $\tau_{decay}$ | $p$ | $delay_{mean}$ | $delay_{\sigma}$ | Compartment |
|---|---|---|---|---|---|---|---|---|
| bis | 1.5 | 0.7 | 0.5 | 4 | 0.2 | 1.2 | 0.2 | dendrite |
| ca3_non_spatial | 0.7 | 0.2 | 2 | 6.3 | 0.03 | 2.5 | 0.5 | dendrite |
| ca3_spatial | 0.7 | 0.2 | 2 | 6.3 | 0.03 | 2.5 | 0.5 | dendrite |
| cckbas | 0.5 | 0.7 | 0.5 | 4 | 0.2 | 1.2 | 0.2 | dendrite |
| ivy | 0.5 | 0.7 | 0.5 | 4 | 0.2 | 1.2 | 0.2 | dendrite |
| mec | 1 | 0.05 | 2 | 6.3 | 0.01 | 10 | 0.5 | dendrite |
| msteevracells | 0.9 | 0.7 | 0.5 | 3 | 0.5 | 10.5 | 0.5 | soma |
| olm | 1.5 | 0.7 | 0.5 | 4 | 0.2 | 1.2 | 0.2 | dendrite |
| pvbas | 1.5 | 0.7 | 0.5 | 4 | 0.2 | 1.2 | 0.2 | dendrite |
| pyr | 0.04 | 0.02 | 0.3 | 0.6 | 0.07 | 1.2 | 0.2 | dendrite |
| sca | 1.5 | 0.7 | 0.5 | 4 | 0.1 | 1.2 | 0.2 | dendrite |
| lec | 0.1 | 0.05 | 2 | 6.3 | 0.003 | 8 | 0.5 | dendrite |

Continued on next page

Table 14: Synaptic connections to bis

| | $g_{max,mean}$ | $g_{max,\sigma}$ | $\tau_{rise}$ | $\tau_{decay}$ | $p$ | $delay_{mean}$ | $delay_{\sigma}$ | Compartment |
|---|---|---|---|---|---|---|---|---|

Table 14: Synaptic connections to bis

| | $g_{max,mean}$ | $g_{max,\sigma}$ | $\tau_{rise}$ | $\tau_{decay}$ | $p$ | $delay_{mean}$ | $delay_{\sigma}$ | Compartment |
|---|---|---|---|---|---|---|---|---|
| bis | 0.5 | 0.2 | 0.5 | 4 | 0.2 | 1.2 | 0.2 | dendrite |
| ca3_non_spatial | 0.02 | 0.01 | 1.3 | 8 | 0.06 | 1.2 | 0.2 | dendrite |
| ca3_spatial | 0.02 | 0.01 | 1.3 | 8 | 0.06 | 1.2 | 0.2 | dendrite |
| cckbas | 0.1 | 0.05 | 0.5 | 4 | 0.2 | 1.2 | 0.2 | dendrite |
| mec | 0.5 | 0.3 | 1.3 | 8 | 0.01 | 10.2 | 0.2 | dendrite |
| pvbas | 0.035 | 0.015 | 0.29 | 2.67 | 0.2 | 1.2 | 0.2 | dendrite |
| pyr | 0.5 | 0.07 | 1.3 | 8 | 0.14 | 1.2 | 0.2 | dendrite |
| sca | 0.1 | 0.05 | 0.5 | 4 | 0.05 | 1.2 | 0.2 | dendrite |

Table 15: Synaptic connections to cckbas

| | $g_{max,mean}$ | $g_{max,\sigma}$ | $\tau_{rise}$ | $\tau_{decay}$ | $p$ | $delay_{mean}$ | $delay_{\sigma}$ | Compartment |
|---|---|---|---|---|---|---|---|---|
| bis | 1 | 0.7 | 0.5 | 4 | 0.2 | 1.2 | 0.2 | dendrite |
| cckbas | 0.2 | 0.2 | 0.2 | 4.2 | 0.63 | 2.7 | 0.5 | dendrite |
| ivy | 0.5 | 0.2 | 0.5 | 4 | 0 | 1.2 | 0.2 | dendrite |
| msteevracells | 3.5 | 0.2 | 0.5 | 5 | 0.5 | 10.5 | 2.5 | dendrite |
| ngf | 0.5 | 0.2 | 0.5 | 10 | 0.1 | 1.2 | 0.2 | dendrite |
| olm | 0.5 | 0.7 | 0.5 | 4 | 0 | 1.2 | 0.2 | dendrite |
| pvbas | 1 | 0.2 | 0.29 | 2.67 | 0.05 | 1.2 | 0.2 | dendrite |

Table 16: Synaptic connections to ivy

| | $g_{max,mean}$ | $g_{max,\sigma}$ | $\tau_{rise}$ | $\tau_{decay}$ | $p$ | $delay_{mean}$ | $delay_{\sigma}$ | Compartment |
|---|---|---|---|---|---|---|---|---|
| cckbas | 0.5 | 0.07 | 0.5 | 4 | 0.1 | 1.2 | 0.2 | dendrite |
| ivy | 0.5 | 0.2 | 0.5 | 4 | 0.5 | 1.2 | 0.2 | dendrite |
| pvbas | 0.5 | 0.2 | 0.5 | 4 | 0.5 | 1.2 | 0.2 | dendrite |
| pyr | 0.041 | 0.021 | 0.3 | 0.6 | 0.13 | 1.2 | 0.2 | dendrite |
| sca | 3.5 | 0.7 | 0.5 | 4 | 0.2 | 1.2 | 0.2 | dendrite |

Table 17: Synaptic connections to ngf

| | $g_{max,mean}$ | $g_{max,\sigma}$ | $\tau_{rise}$ | $\tau_{decay}$ | $p$ | $delay_{mean}$ | $delay_{\sigma}$ | Compartment |
|---|---|---|---|---|---|---|---|---|
| ca3_non_spatial | 1.42 | 0.6 | 0.5 | 3 | 0.2 | 1.5 | 0.5 | dendrite |

Continued on next page

Table 17: Synaptic connections to ngf

| | $g_{max,mean}$ | $g_{max,\sigma}$ | $\tau_{rise}$ | $\tau_{decay}$ | $p$ | $delay_{mean}$ | $delay_{\sigma}$ | Compartment |
|---|---|---|---|---|---|---|---|---|
| ca3_spatial | 1.42 | 0.6 | 0.5 | 3 | 0.2 | 1.5 | 0.5 | dendrite |
| ivy | 0.9 | 0.7 | 3.1 | 42 | 0.8 | 1.2 | 0.2 | dendrite |
| mec | 1.7 | 0.3 | 0.5 | 3 | 0.3 | 10.2 | 0.2 | dendrite |
| ngf | 0.75 | 0.7 | 3.1 | 42 | 0.7 | 1.2 | 0.2 | dendrite |
| olm | 1.27 | 0.6 | 1.3 | 10.2 | 0.4 | 1.2 | 0.2 | dendrite |
| lec | 0.7 | 0.3 | 0.5 | 3 | 0.05 | 1.2 | 0.2 | dendrite |

Table 18: Synaptic connections to olm

| | $g_{max,mean}$ | $g_{max,\sigma}$ | $\tau_{rise}$ | $\tau_{decay}$ | $p$ | $delay_{mean}$ | $delay_{\sigma}$ | Compartment |
|---|---|---|---|---|---|---|---|---|
| ivy | 1.5 | 0.7 | 0.5 | 4 | 0.5 | 1.2 | 0.2 | dendrite |
| msach | 1.5 | 0.1 | 0.5 | 3 | 0.6 | 10.5 | 0.5 | soma |
| pyr | 0.031 | 0.0015 | 0.3 | 0.6 | 0.081 | 1.2 | 0.2 | dendrite |
| sca | 1.5 | 0.7 | 0.5 | 4 | 0.2 | 1.2 | 0.2 | dendrite |

Table 19: Synaptic connections to pvbas

| | $g_{max,mean}$ | $g_{max,\sigma}$ | $\tau_{rise}$ | $\tau_{decay}$ | $p$ | $delay_{mean}$ | $delay_{\sigma}$ | Compartment |
|---|---|---|---|---|---|---|---|---|
| bis | 1.1 | 0.05 | 0.5 | 4 | 0.5 | 1.2 | 0.2 | dendrite |
| ca3_non_spatial | 0.9 | 0.1 | 2 | 6.3 | 0.06 | 1.5 | 0.5 | dendrite |
| ca3_spatial | 750 | 0.2 | 2 | 6.3 | 0.6 | 1.5 | 0.5 | dendrite |
| cckbas | 10 | 0.2 | 0.43 | 4.49 | 0.38 | 4.5 | 2 | soma |
| ivy | 10.1 | 0.5 | 0.5 | 4 | 0.8 | 1.2 | 0.2 | dendrite |
| ngf | 1 | 0.05 | 0.5 | 4 | 0.8 | 1.2 | 0.2 | dendrite |
| olm | 0.73 | 0.35 | 0.25 | 7.5 | 0.5 | 1.2 | 0.2 | dendrite |
| pvbas | 10 | 0.01 | 0.8 | 4.8 | 0.7 | 1.2 | 0.2 | soma |
| pyr | 0.5 | 0.04 | 0.07 | 0.2 | 0.13 | 1.2 | 0.2 | dendrite |
| sca | 1.1 | 0.05 | 0.5 | 4 | 0.5 | 1.2 | 0.2 | dendrite |

Table 20: Synaptic connections to pyr

| | $g_{max,mean}$ | $g_{max,\sigma}$ | $\tau_{rise}$ | $\tau_{decay}$ | $p$ | $delay_{mean}$ | $delay_{\sigma}$ | Compartment |
|---|---|---|---|---|---|---|---|---|
| aac | 20.5 | 10 | 0.28 | 8.4 | 0.29 | 1.2 | 0.2 | axon |
| bis | 0.009 | 0.005 | 0.11 | 9.7 | 0.14 | 1.2 | 0.2 | dendrite |
| ca3_non_spatial | 0.016 | 0.002 | 0.5 | 3 | 0.4 | 1.5 | 0.5 | rad |
| ca3_spatial | 3000 | 0.9 | 0.5 | 3 | 0.6 | 1.5 | 0.5 | rad |
| cckbas | 2.5 | 1.2 | 0.2 | 4.2 | 0.63 | 2.5 | 1.2 | soma |
| ivy | 0.053 | 0.02 | 1.1 | 11 | 0.13 | 1.2 | 0.2 | lm |

Continued on next page

Table 20: Synaptic connections to pyr

| | $g_{max,mean}$ | $g_{max,\sigma}$ | $\tau_{rise}$ | $\tau_{decay}$ | $p$ | $delay_{mean}$ | $delay_{\sigma}$ | Compartment |
|---|---|---|---|---|---|---|---|---|
| mec | 1500 | 0.4 | 0.5 | 3 | 0.6 | 10 | 2 | lm |
| ngf | 0.098 | 0.05 | 9 | 39 | 0.29 | 1.2 | 0.2 | lm |
| olm | 1.7 | 0.9 | 0.13 | 11 | 0.29 | 1.2 | 0.2 | lm |
| pvbas | 500 | 0.9 | 0.3 | 6.2 | 0.8 | 1.2 | 0.2 | soma |
| pyr | 0.01 | 0.007 | 0.1 | 1.5 | 0.01 | 1.2 | 0.2 | basal |
| sca | 0.098 | 0.05 | 0.3 | 6.2 | 0.29 | 1.2 | 0.2 | rad |
| lec | 0.06 | 0.007 | 0.5 | 3 | 0.07 | 10 | 2 | lm |

Table 21: Synaptic connections to sca

| | $g_{max,mean}$ | $g_{max,\sigma}$ | $\tau_{rise}$ | $\tau_{decay}$ | $p$ | $delay_{mean}$ | $delay_{\sigma}$ | Compartment |
|---|---|---|---|---|---|---|---|---|
| bis | 0.5 | 0.2 | 0.5 | 4 | 0.5 | 1.2 | 0.2 | dendrite |
| ca3_non_spatial | 0.05 | 0.02 | 0.5 | 4 | 0.06 | 1.2 | 0.2 | dendrite |
| ca3_spatial | 0.05 | 0.02 | 0.5 | 4 | 0.06 | 1.2 | 0.2 | dendrite |
| ivy | 0.5 | 0.2 | 0.5 | 4 | 0.1 | 1.2 | 0.2 | dendrite |
| ngf | 0.5 | 0.2 | 0.5 | 4 | 0.5 | 1.2 | 0.2 | dendrite |
| olm | 1.3 | 0.6 | 0.07 | 29 | 0.1 | 1.2 | 0.2 | dendrite |
| sca | 0.03 | 0.015 | 4 | 34.3 | 0.3 | 1.2 | 0.2 | dendrite |

The weights of connections from the ca3_spatial and mec generators and PV basket neurons to pyramidal cells and from pyramidal neurons and ca3_spatial generators to PV basket neurons were computed by the Gaussian function

$$g = g_{max,conn} exp\Big( - 0.5 \cdot \frac{D_{conn}^2}{\sigma_{conn}^2} \Big) \cdot \frac{S_{conn}}{\sigma_{conn} \cdot \sqrt{2\pi}} \qquad (165)$$

Table 22: Parameters of spatial modulated connection

| Connection type | $\sigma_{conn}$ | $S_{conn}$ | $D_{conn}$ |
|---|---|---|---|
| ca3_spatial → pyr | 300 | 37500 | $x_{pre} - x_{post}$ |
| mec → pyr | 300 | 37500 | $x_{pre} + 500 - x_{post}$ |
| pvbas → pyr | 300 | 500 | $1/(x_{pre} - x_{post})$ |
| pyr → pvbas | 27000 | 0.5 | $x_{pre} - x_{post}$ |
| ca3_spatial → pvbas | 27000 | 750 | $x_{pre} - x_{post}$ |

where $x_{pre}$ and $x_{post}$ are the coordinates of pre-and postsynaptic neurons in the connections space. The coordinates for the pyramidal and PV basket neurons were distributed linearly in increments of 3 ms and 50 ms, respectively. Half of the pyramid neurons did not have an x coordinate, and the connections of this population were set randomly, as for all other populations of neurons. For the ca3_spatial generators, $x$ was equal to their peak firing rate. For the mec generators, $x$ is the peak of the discharges closest to the coordinate of the pyramid neuron, the peaks of the mec discharges are calculated based on the frequency and phase of the grid.

### 6 Gap junction models

#### 6.1 General description of gap junction models

Gap junction between two neurons (1 and 2) is described by a simple equation:

$$I_{gap} = \frac{V_1 - V_2}{R_{gap}} \tag{166}$$

where $I_{gap}$is current for neuron 1, $V_1$ and $V_2$ are potentials of neuron 1 and 2 respectively. Gap junctions are always symmetrical, in the equation of neuron 2 the same current is added. $R_{gap}$ is the resistance of contact, it is chosen randomly from a normal distribution for each contact, parameters see below.

Table 23: Parameters of gap junctions

| Celltype | $R_{gap,mean}$ | $R_{gap,\sigma}$ | $p$ | Compartment |
|---|---|---|---|---|
| ngf | 1000000.0 | 10 | 0.7 | dendrite |
| pvbas | 100000.0 | 10 | 0.1 | dendrite |

### 7 Simulation of local field potetial

LFP is simulated with LFPsim tools [4], we have used line approximation:

$$\Phi_{LFP} = \sum_{i=1}^{n} \frac{I_i}{2\pi\sigma} log\Big(\frac{\sqrt{h_i^2 + r_i^2} - h_i}{\sqrt{l_i^2 + r_i^2} - l_i}\Big) \tag{167}$$

$\Phi_{LFP}$ is simulated LFP, $I$ is transmembrane current. In the simulation of LFP, only compartments of pyramidal neurons were taken into account. $\sigma$ denotes the conductivity of the medium. $r$ is the distance from the source to the point of measurement, $h$ is the distance from the end of a segment to point of measurement, $l$ is the length of a segment. All the pyramid neurons were arranged in an orderly fashion. TAll the pyramid neurons were arranged in an orderly fashion. The soma of all the cells was on the same level ($z = 0$). The x and y coordinates for each neuron were chosen randomly from the circle with radius $R_{x,y}$ specified by the formula:

$$R_{x,y} = \frac{\sqrt{\frac{N_{pyr}}{\rho_{pyr}}}}{\pi} \tag{168}$$

where $N_{pyr}$ is the number of pyramidal cells in the model, $\rho_{pyr} = 0.2\ cell/\mu m^2$ is the density of cells in the pyramidal layer of the CA1 field.

[3] Directional statistics. URL http://doi.wiley.com/10.1002/9780470316979.

[4] Harilal Parasuram, Bipin Nair, Egidio D'Angelo, Michael Hines, Giovanni Naldi, and Shyam Diwakar. Computational modeling of single neuron extracellular electric potentials and network local field potentials using LFPsim. 10. ISSN 1662-5188. doi: 10.3389/fncom.2016.00065. URL https://www.frontiersin.org/articles/10.3389/fncom.2016.00065/full. Publisher: Frontiers.

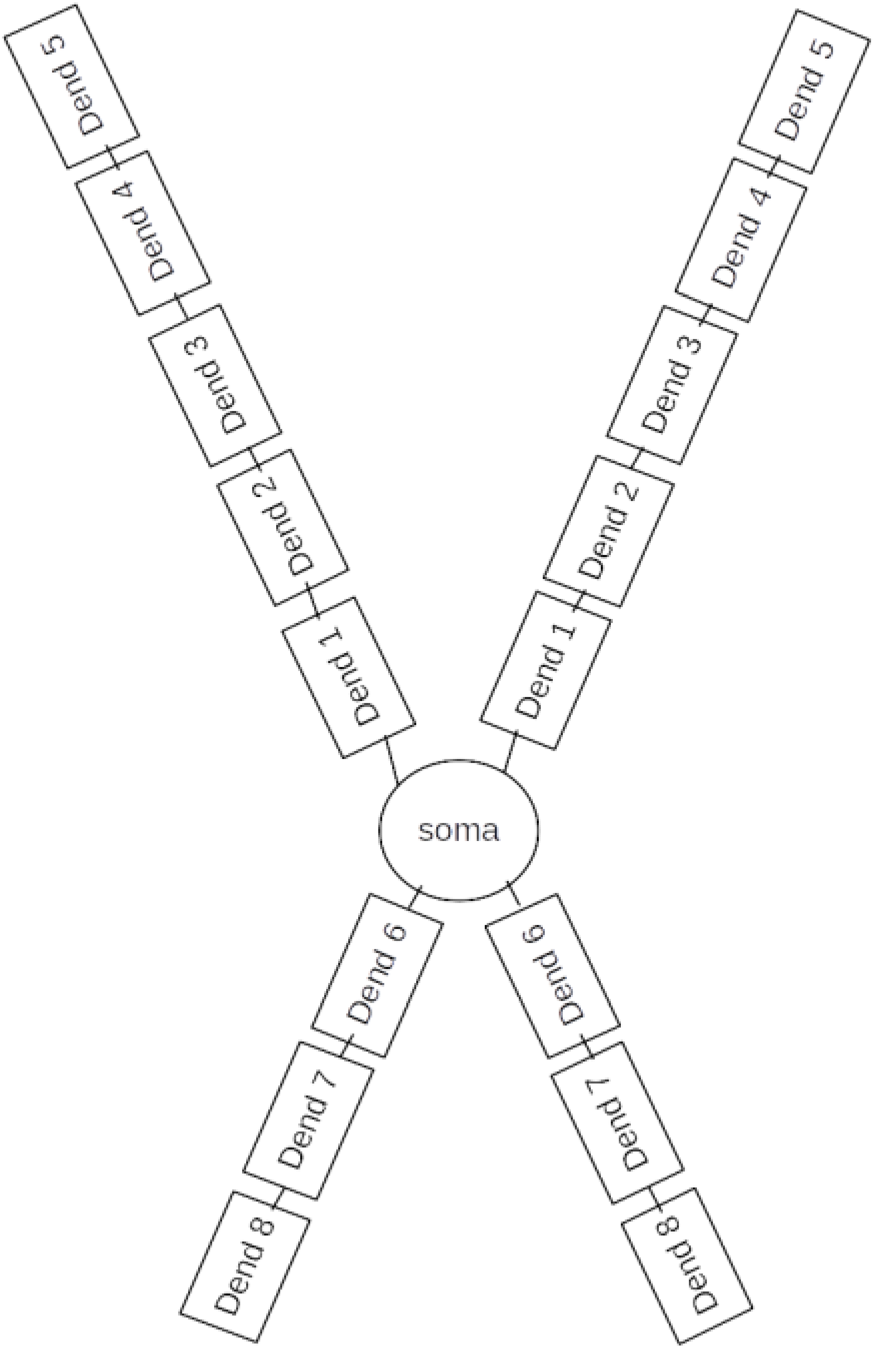


Figure 1: Scheme of compartments of sca, ivy, ngf, pvbas, cckbas, aac cells

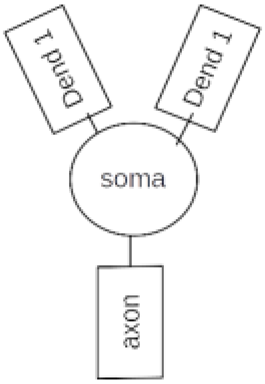


Figure 2: Scheme of compartments of olm cell

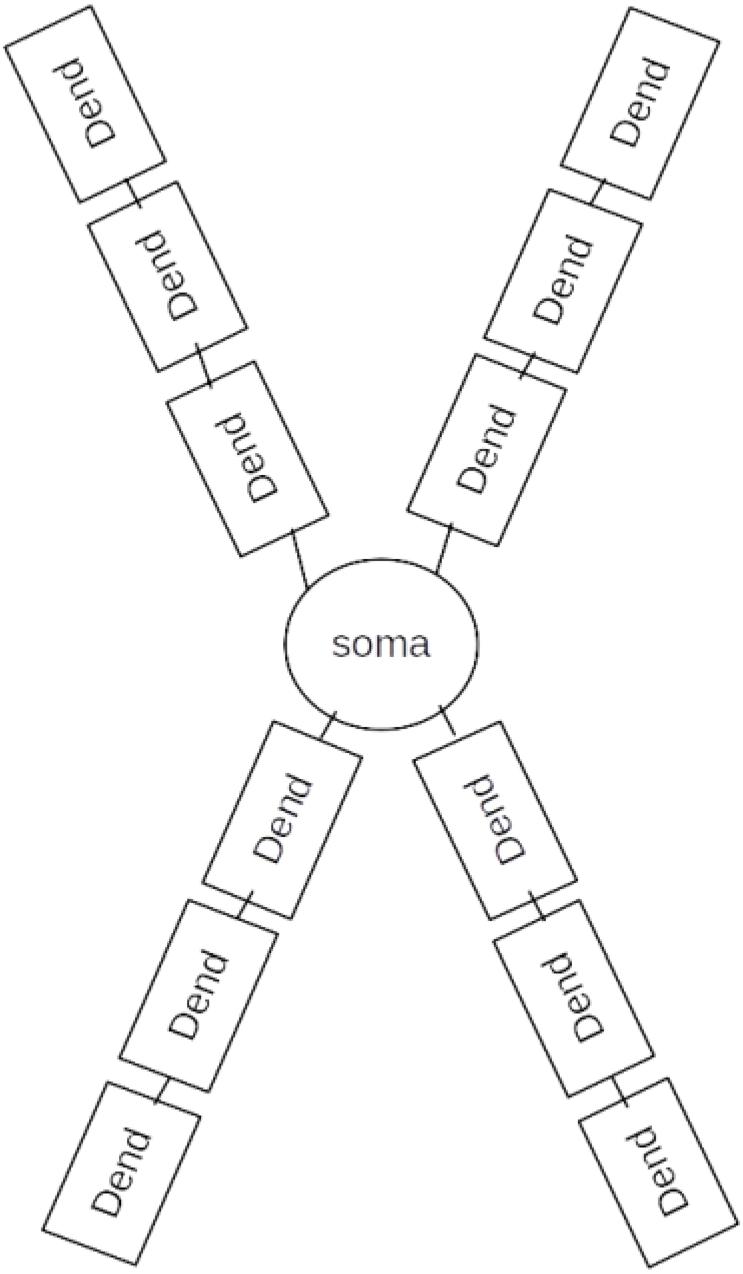


Figure 3: Scheme of compartments of bis cell

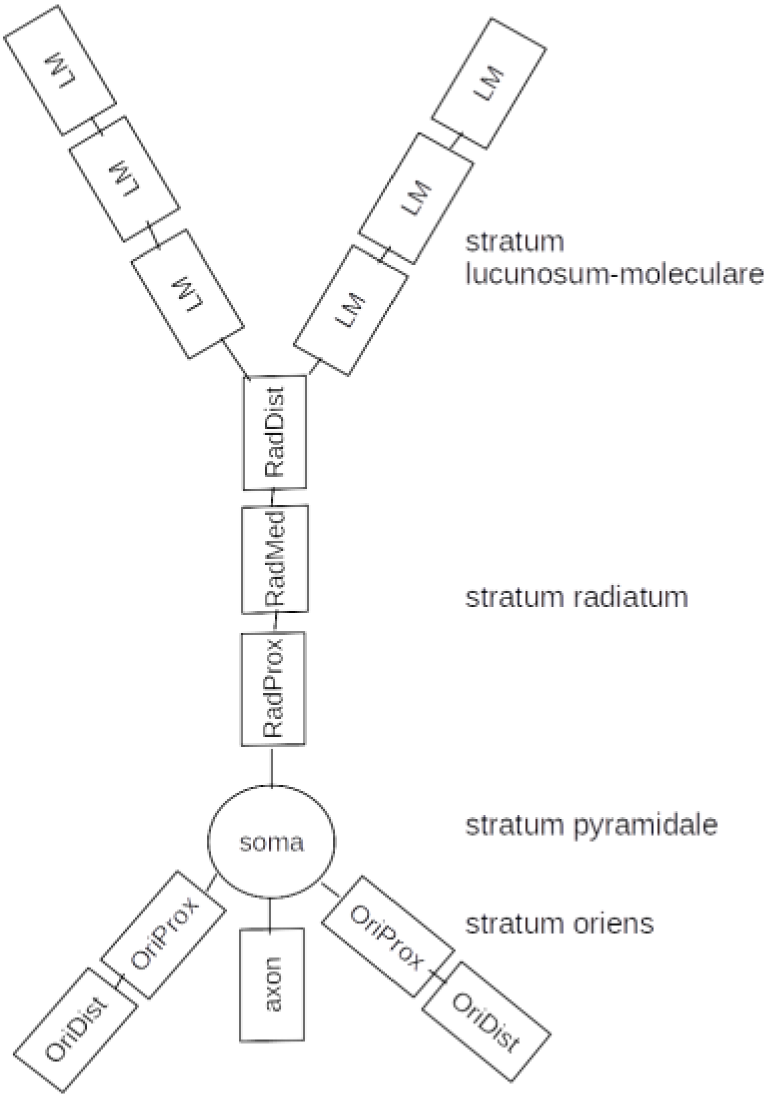


Figure 4: Scheme of compartments of pyramidal cell
